# Septins buffer actomyosin forces to protect the nucleus from genotoxic mechanical stress

**DOI:** 10.64898/2026.01.21.700789

**Authors:** Franklin Mayca-Pozo, Sarah M. Butts, Andrew W. Schaefer, Cristina Montagna, Elias T. Spiliotis

## Abstract

Invasively migrating cells thread their nucleus through confined interstitial spaces. How cells protect the nucleus from intracellular forces is poorly understood. Here, we show that the septin cytoskeleton buffers the actomyosin forces that power nuclear movement. Septin filaments comprising SEPT9, a septin amplified in breast cancer, align with perinuclear actomyosin cables which exhibit higher tensile stress during 3D confined migration through narrower pores. SEPT9 depletion amplifies actin stress during confined migration and after myosin II hyper-activation in non-migrating cells causing actin and nuclear envelope ruptures. Following confined migration, DNA breaks, nuclear blebs, micronuclei and cell death increase in SEPT9-depleted cells, phenotypes rescued by the oncogenic SEPT9 isoform 1. Clinicogenomic data reveal that *SEPT9* amplification associates with lower genomic alteration in aggressive breast tumors and higher patient mortality. We propose that septins are a mechanoprotective element of the cytoskeleton, and SEPT9 amplification enhances tumor cell survival by preventing nuclear damage.

## Main

Cell migration is a fundamental biological process underlying organismal development, tissue homeostasis and regeneration, immunity, and cancer metastasis.^1,2^ In the three-dimensional environment of tissues and interstitial spaces, cell migration is physically constrained by the dimensions of the surrounding microenvironment.^3^ The nucleus, the largest and most rigid organelle in the cell, poses a major physical impediment that restricts cell mobility.^4,5^ Cells overcome this limitation by modifying the physical properties of the nucleus and/or the extracellular matrix (ECM),^6–9^ and by employing different strategies and modes of migration.^10–12^

The nucleus is a mechanosensitive organelle, sensing not only external spatial constraints,^13–15^ but also forces transmitted from the ECM through the cytoskeleton.^5,8,9,16^ Nuclear mechanosensitivity is a central determinant of cell behavior and identity, as gene expression and nucleocytoplasmic transport are directly impacted by forces acting on chromatin and nuclear pores.^17–19^ While advantageous for cellular homeostasis, nuclear mechanosensitivity poses a risk to nuclear and genome integrity. During confined migration, the nucleus is subjected to multiple forces (e.g, contractile, frictional, hydrostatic), which collectively impose mechanical stresses capable of rupturing the nuclear envelope and damaging genomic DNA.^20,21^ These injuries alter nucleocytoplasmic transport and risk DNA leakage into the cytoplasm, potentially leading to further genomic alterations, cell death, or the pathogenesis of diseases such as autoimmunity and cancer.^22–24^

How cells mitigate mechanical stress to avoid nuclear damage during confined migration is not well understood. In the nucleoplasm, lamin intermediate filaments provide structural support and determine nuclear stiffness, dictating how nuclei respond to mechanical stress and deformation.^6,25,26^ Softer nuclei with lower levels of lamin A/C are more amenable to confined migration but are also susceptible to nuclear damage, which may be compensated by as-yet unidentified factors.^27–29^ In the cytoplasm, vimentin filaments form a cage-like structure around the nucleus, protecting it from rupture by reinforcing perinuclear stiffness while absorbing contractile forces.^30^ Recent work suggests that vimentin functions in conjunction with perinuclear microtubules to buffer compressive forces.^31,32^ However, vimentin also promotes the tensile forces of cell migration, which are powered by actomyosin filaments anchored to focal adhesions.^33,34^ Vimentin’s dual role in absorbing and transmitting forces, along with its context-dependent effects as either an inhibitor or enhancer of cell migration, points to additional mechanisms of force regulation that remain to be elucidated.

Septins comprise a distinct network of non-polar cytoskeletal polymers that assemble from multimeric GTP-binding proteins.^35,36^ Septins associate with subsets of actin filaments and microtubules, as well as specific membrane subdomains, regulating the spatial organization of cellular proteins and processes.^37–39^ Septin 9 (SEPT9) is ubiquitously expressed and mediates the interaction of septin oligomers and polymers with actin and microtubules.^40–44^ SEPT9 isoform 1 (SEPT9_i1) is the longest SEPT9 isoform and possesses actin-binding and crosslinking activity.^41,44,45^ SEPT9_i1 has been functionally implicated in angiogenesis,^46^ cell migration^41,47–50^ and ECM degradation by invadosomes.^51,52^ Notably, overexpression and depletion of SEPT9_i1 enhances and reduces, respectively, tumor growth and metastasis in vivo.^48,53,54^

The mechanical properties of septins in cell migration and broader cell biology remain largely unexplored. Recent work indicated that SEPT9_i1 influences nuclear deformability, impacting the formation of invadopodia,^52^ which occurs partly in response to the inability of the nucleus to overcome physical confinement,^55,56^ and may involve a mechanical crosstalk between the nuclear and plasma membrane ends of invadopodia.^57^ However, whether septins function in nuclear mechanosensation and/or mechanotransduction during confined cell migration remains unknown. Here, we report that septins are integral components of the force-bearing actomyosin cytoskeleton that surrounds the pore-confined nucleus, where they buffer the contractile forces that drive nuclear transit. We demonstrate that SEPT9_i1 protects the nucleus from genotoxic mechanical stress and enhances cell survival by reducing genomic damage. Clinicogenomic analyses reveal that *SEPT9* copy number amplification associates with distinct thresholds of genomic alteration in aggressive breast cancer tumors and correlates with reduced patient survival. We propose that *SEPT9* amplification favors the survival of cancer cells with elevated genomic instability by buffering mechanical stress during confined migration, thereby preventing genomic catastrophe and facilitating metastatic progression.

## Results

### Septin filaments align with contractile actomyosin cables in cellular regions of confined nuclear threading

We sought to investigate how septins function during confined nuclear movement. We used a 3D assay of cell migration through micron scale (3-8 μm wide) pores of track-etched polymeric membranes, which were coated with ECM (Matrigel/laminin) (Fig. **1a**). Bathed in media from both apical and basal sides, the 10-15 μm-deep pores serve as interstitial-like conduits of chemotactic migration along a concentration gradient of epidermal growth factor (EGF). We combined this biomimetic platform with 3D super-resolution SoRa spinning disk confocal microscopy (SDCM), which provided an unprecedented view and resolution of the cellular regions confined within the micron-scale environment of ECM-containing pores.

**Fig. 1:**
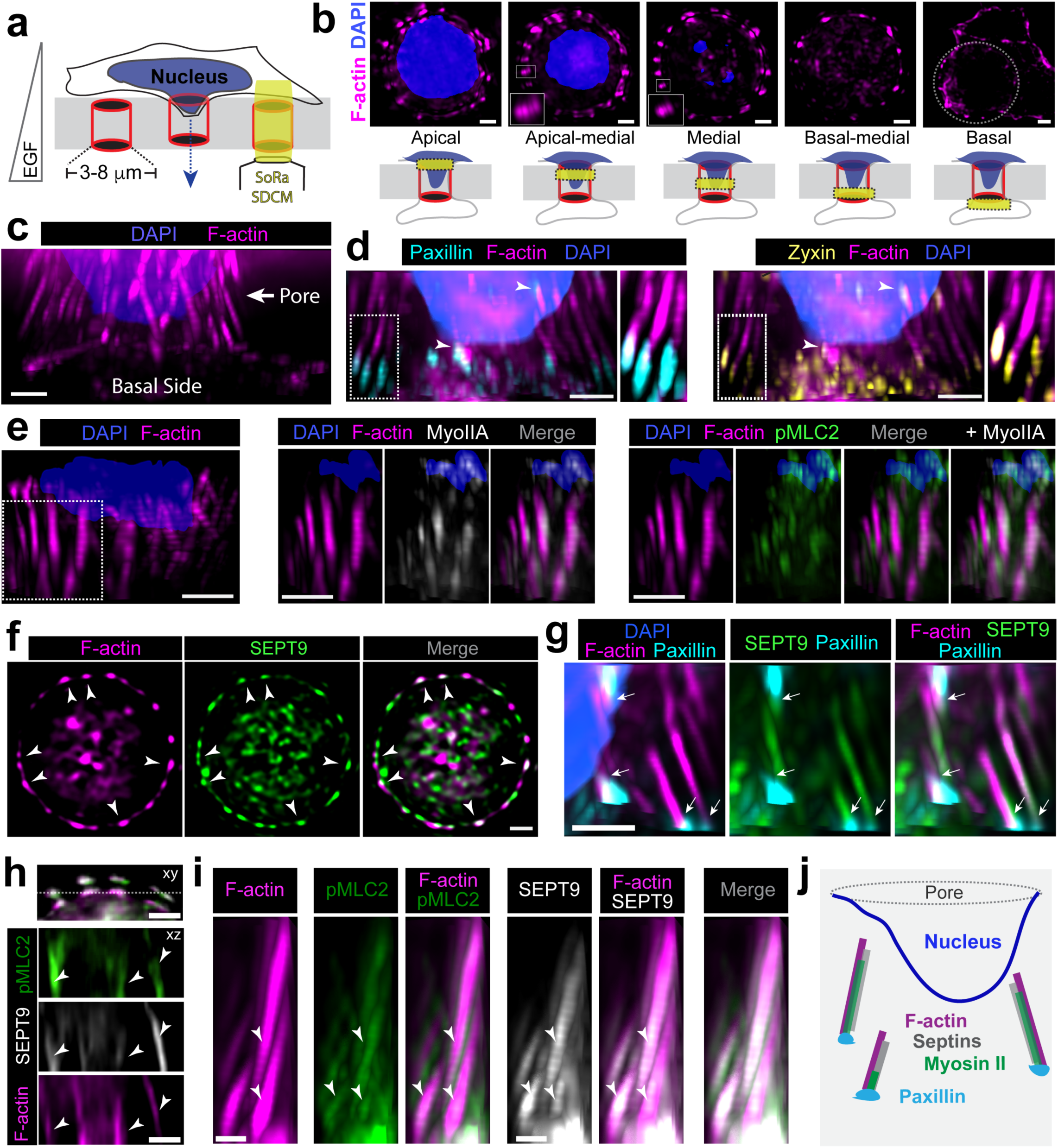
Septins align with perinuclear contractile actomyosin cables during confined cell migration. **a**, Schematic of the experimental configuration for imaging confined cell migration in 3D. Transwell polycarbonate membranes with pores of variable diameter were coated with ECM, and U2OS cells migrated through the pores by chemotaxis along an EGF gradient. Cellular regions within the pore were imaged by super-resolution SoRa SDCM. **b,** Maximum intensity projections of cross-sectional images from ∼1 μm-thick regions at apical, apical-medial, medial, basal-medial, and basal depths of the pore, indicated by the yellow band in the schematic below each fluorescence image. Insets show magnified views of phalloidin-stained F-actin clusters (groups of vertical filaments; magenta) surrounding the confined nuclear mass (DAPI, blue) with semi-regular spacing along the pore circumference. The basal cross-section shows cortical actin at the cell membrane extending from the pore (dotted circle) to the basal surface of the porous membrane. **c,** Side view of a 3D volume showing F-actin (phalloidin, magenta) relative to the confined nucleus (DAPI, blue) within the pore and the plasma membrane extending onto the basal surface. **d,** Sagittal view of 3D-rendered images of the confined nucleus (DAPI, blue), F-actin (phalloidin, magenta), paxillin (cyan), and zyxin (yellow). **e,** Sagittal views of 3D-rendered images showing myosin IIA (grayscale) and pMLC2 (green) localization to F-actin cables (phalloidin, magenta) in the perinuclear region outlined in the leftmost panel (dashed rectangle). **f,** Mediobasal optical section showing F-actin (phalloidin, magenta) and SEPT9 (green) colocalization (arrowheads) along the pore circumference. **g,** Sagittal view of perinuclear actin cables (phalloidin, magenta) colocalizing with SEPT9 (green) and contacting paxillin (cyan; arrows). **h,** SoRa SDCM images show single optical xy (top panel) and xz sections across the plane outlined with a dashed line. In top panel, F-actin (phalloidin, magenta), SEPT9 (grayscale) and pMLC2 (green) are merged. Arrowheads point to actin segments with pMLC2 and SEPT9. **i,** Side view of a subcellular region stained for F-actin (phalloidin, magenta), pMLC2 (green) and SEPT9 (grayscale) after 3D rendering. Arrowheads point to actin segments with pMLC2 and SEPT9. Scale bars, 0.5 μm. **j,** Schematic illustrates the organization of perinuclear septoactomyosin cables in regions of confined nuclear movement. All U2OS cell images were acquired with super-resolution SoRa SDCM 12 h after plating on 8 μm-wide pores. In 3D views, occasional banding of F-actin fluorescence was due to sub-Nyquist sampling, which was necessary to offset photobleaching during acquisition of large image z-stacks. Scale bars, 1 μm.

Imaging the actin cytoskeleton of the human osteosarcoma U2OS cells revealed an arrangement of actin cables that extend vertically in the apicobasal direction of cell migration, surrounding the constricted portion of the nucleus that has entered the pore (Fig. **1b**). These cables were oriented parallel to the confined nuclear mass without forming discernible contacts with each other or the nucleus, resembling the vertical bars of a birdcage (Fig. **1c**, **Supplementary Video 1**). Transverse optical sections taken from the apex to the base of the pore, together with 3D image reconstructions, showed that actin cables initiate and terminate at different pore depths, with several cables overlapping vertically one above the other in stacked groups of two, three or more (see insets in Fig. **1b,d**, **Supplementary Video 1**). In 8 μm-wide pores, actin cables displayed a circular arrangement with semi-regular spacing (Fig. **1b**), though this symmetrical organization was not as consistent in narrower pores.

To determine whether the perinuclear actin cables transduce forces, we examined the localization of force-sensing proteins paxillin and zyxin, which accumulate at focal adhesions and sites of actin filament strain,^58,59^ myosin II heavy chain (myoIIA), and the phosphorylated (pT18/pS19) myosin regulatory light chain (pMLC2), as a marker for myosin II contractility.^60^ Actin cables were anchored to focal concentrations of paxillin and zyxin, which localized predominantly on the basal side of the pore distal to the edge of the confined nucleus (Fig. **1d**, insets; **Supplementary Video 1**).

Occasionally, actin cables contacted paxillin and zyxin foci in closer proximity to the nucleus at apical and medial regions of the pore (Fig. **1d**, arrowheads). Consistent with active force transduction, actin cables were closely juxtaposed to myoIIA, which appeared as thick cable-like filaments similar to actin, as well as pMLC2, which localized to patches and segments along the actomyosin cables (Fig. **1e**). Together, these findings suggest that the nucleus is surrounded by contractile actomyosin cables that generate forces that may promote nuclear movement in the apicobasal direction of migration.

We next examined the localization of septin filaments and vimentin intermediate filaments, which associate with and function synergistically with actomyosin during cell migration.^33,34,37^ We immunostained for SEPT9 and SEPT7, which are ubiquitous and requisite subunits of the octameric septin protomer.^36^ Using SEPT9 pan-isoform and SEPT9_i1 isoform-specific antibodies, we found that septin filaments exhibited similar localization and distribution to perinuclear actomyosin cables (Fig. **1f** and **Extended Data Fig. 1a**; **Supplementary Video 2**). SEPT9 filaments were closely juxtaposed to actin cables, overlapping laterally along their vertical length (Fig. **1g****; Supplementary Video 2**). SEPT9 was present on actin cables that connected to paxillin foci (Fig. **1g**, arrows) and contained pMCL2 (Fig. **1h,i**; arrowheads; **Supplementary Video 3**), indicating that septins are an integral component of the actomyosin mechanotransduction machinery (Fig. **1j**). The percentage of actin cables per cell in contact with paxillin was 90% ± 8%, and 84% ± 4% of these were decorated with SEPT9 (*n* = 8 cells).

Consistent with septin filaments composed of the canonical SEPT2/6/7/9 protomer, SEPT7 exhibited similar localization to SEPT9 colocalizing with the oncogenic SEPT9_i1 isoform (**Extended Data Fig. 1b-d**). In contrast, vimentin differed markedly in its localization from the septoactomyosin cables. Vimentin intermediate filaments accumulated predominantly on the basal side of the nucleus, forming a basket-like organization around the ventral nuclear edge and a dense meshwork along the ventral plasma membrane emerging from the basal side of the pore (**Extended Data Fig. 1e**). Perinuclear vimentin filaments with an organization resembling the vertical septoactomyosin cables were scarce and seldomly observed.

To gain insight into the spatiotemporal dynamics of actin relative to the confined nucleus and of septins with respect to actin in live cells, we performed SoRa SDCM of the nuclear membrane labeled with LAP2β-AcGFP1, GFP-SEPT9_i1, and the cell-permeable actin-imaging probe SPY650-FastAct_X. We overcame the optical constraints of imaging living cells in the pores of membrane inserts by cutting out and mounting a membrane strip on a glass-bottom dish, where it was immobilized using a four-well silicon insert with an adhesive biocompatible surface. Super-resolution 4D imaging (3D time-lapse) resolved the nuclear membrane (LAP2β-GFP) as a ring wrapping around the inner perimeter of the pore along its depth (**Extended Data Fig. 2a; Supplementary Video 4**).

Indicative of the forces exerted on the nucleus during confined migration, nuclear membrane folds and creases occurred intermittently (**Extended Data Fig. 2a**, arrows**; Supplementary Video 4**). In 2D optical slices and top-down views, actin filaments appeared as puncta studding the nuclear membrane perimeter (**Extended Data Fig. 2a**). In 3D volume views, these resembled the vertical phalloidin-labeled cables observed in fixed cells (Fig. **1b-c**).

Actin cables were rather stable - lifetimes often exceeded one hour – but they exhibited heterogeneous dynamics (**Extended Data, Fig. 1b-c**). Some actin cables remained spatiotemporally stationary (**Extended Data Fig. 2b-c**, white arrows), while others grew in length and width after forming de novo (**Extended Data Fig. 2b-c**, blue arrowheads) or disassembled (**Extended Data Fig. 2c**, yellow arrowheads). Consistent with our findings in fixed cells, GFP-SEPT9_i1 colocalized with actin cables and displayed similar dynamics (**Extended Data Fig. 2b**, **Supplementary Video 5**). Collectively, these results reveal that during confined migration, septin filaments align with contractile actomyosin cables in a dynamic birdcage-like perinuclear arrangement that is distinct from the vimentin cytoskeleton.

### Septins buffer actomyosin forces, protecting actin cables from increasing mechanical strain in narrower pores

Given that actomyosin cables surrounding the confined nucleus exhibit active mechanotransduction, we investigated how they respond to mechanical strain and how septins function in this process. We reasoned that if actomyosin cables actively drive nuclear threading, they would experience increasing mechanical tension in narrower pores, as higher forces might be needed to move the nucleus forward – due to greater physical constraint and possibly fewer actomyosin cables owing to reduced cytoplasmic volume. To test this prediction, we decreased stepwise the diameter size of Matrigel-filled pores and imaged zyxin in U2OS cell regions confined in 8 μm, 5 μm, and 3 μm wide pores. We also generated stable clones of SEPT9_i1-depleted U2OS cells using CRISPR/Cas9 to assess whether the septins and more specifically SEPT9_i1 impact the mechanical strain of confined migration (**Extended Data Fig. 3a-b**).

In optical sections from the apical, medial, and basal regions of the pore, zyxin levels appeared to increase with decreasing pore diameters (Fig. **2a-c**, **Extended Data Fig. 3c-e**). We quantitatively assessed whether actin cables were subjected to higher mechanical strain by measuring the percentage of zyxin-decorated actin cables per cell in 3D-rendered image volumes (Fig. **2d**, arrows). To distinguish mechanical strain along actin cables from tension at focal adhesions, we excluded actin cables with zyxin localized solely at their ends. The percentage of zyxin-decorated actin cables increased markedly in 3 μm compared to 8 μm wide pores (Fig. **2e**). Consistent with enhanced actomyosin activity, the percentage of actin cables decorated with pMLC2 also increased in 3 μm compared to 8 μm wide pores (Fig. **2f**). Notably, SEPT9 presence on actin cables was similarly elevated in the narrower 3 μm wide pores (Fig. **2g**), suggesting that SEPT9-actin association responds to the mechanical forces of confined migration. Strikingly, SEPT9_i1 depletion resulted in a substantial increase of zyxin levels throughout the confined perinuclear cytoplasm in pores of all diameter sizes (Fig. **2a-c**, **e**; **Extended Data Fig. 3c-e**).

**Fig. 2:**
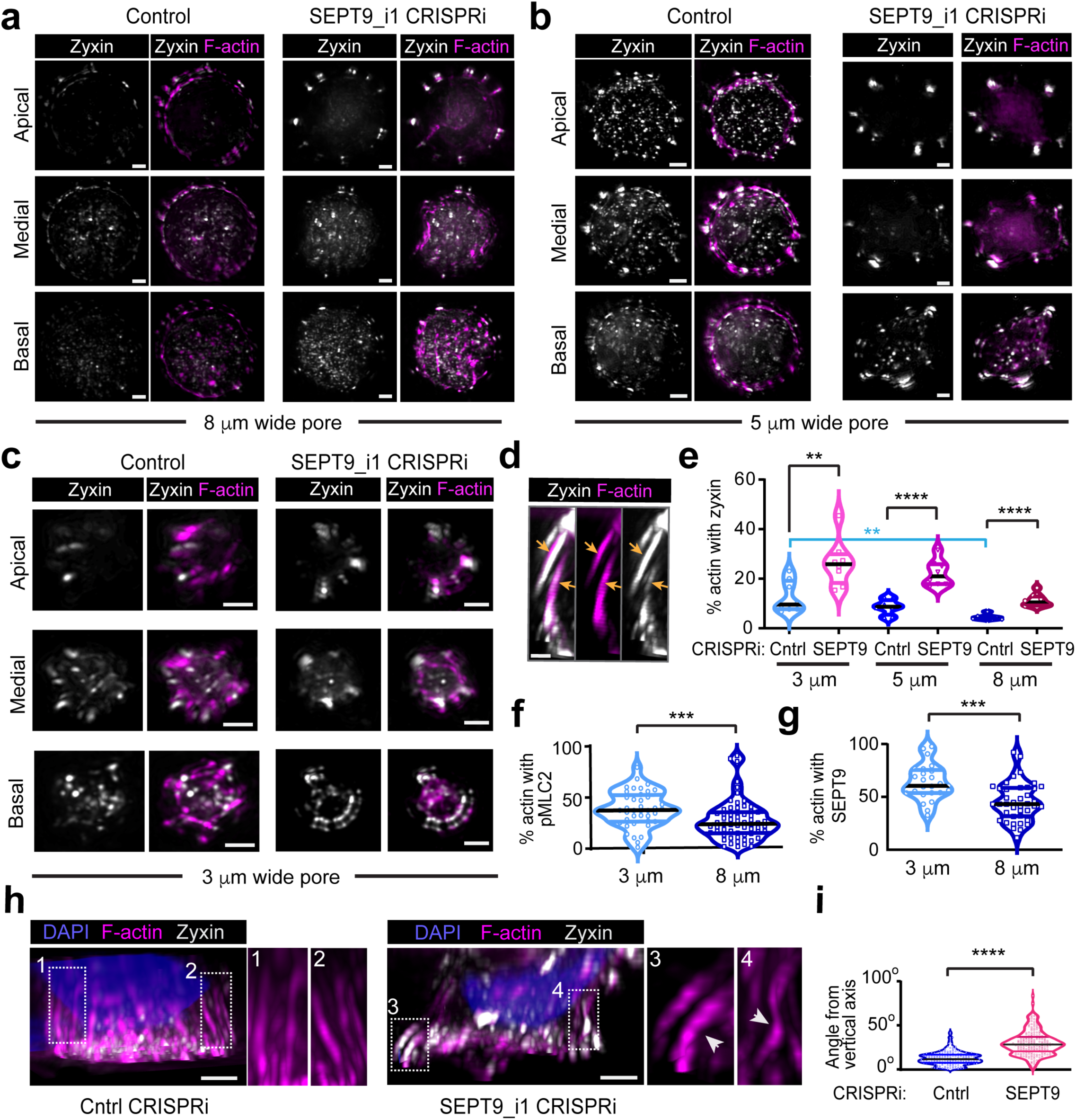
Septins reduce mechanical stress on the actomyosin cytoskeleton during migration through increasingly narrow pores. **a-c**, Maximum intensity projections of cross-sectional images from ∼1 μm-thick regions at apical, medial and basal depths of 8 μm (a), 5 μm (b) and 3 μm (c) wide pores. The confined regions of genome-edited U2OS cells (parental control and SEPT9_i1 CRISPRi) are shown after staining for F-actin (phalloidin, magenta) and zyxin (grayscale). Scale bars, 1 μm. **d,** Sagittal view of a subcellular 3D volume region of a SEPT9_i1-depleted U2OS cell confined in a 5 μm wide pore. Arrows point to zyxin (grayscale) localization along the length of actomyosin cables (phalloidin, magenta). Scale bar, 0.5 μm. **e,** Violin plots show percentage of zyxin decorated actin cables in genome-edited U2OS cell (parental control and SEPT9_i1 CRISPRi) regions confined in 3 μm (*n* = 96-102 cables; 10 cells), 5 μm (*n* = 185-190 cables; 10 cells) and 8 μm (*n* = 381-390 cables; 10 cells) pores. Statistical analysis between the indicated pair groups was done with the Welch’s t-test. **f,** Violin plot shows pMLC2-decorated actin cables with SEPT9 per cell regions confined in 3 μm (*n* = 40) and 8 μm (*n* = 63) wide pores. Statistical analysis was done with a Mann Whitney U-test. ***, p < 0.001. **g,** Violin plot shows percentage of SEPT9-decorated actin cables per cell region confined in 3 μm (*n* = 25) and 8 μm (*n* = 40) wide pores. Statistical analysis was done with Welch’s t-test. **h,** Sagittal view of 3D volume regions of control and SEPT9_i1 depleted U2OS cells confined in 8 μm wide pores after staining for actin (phalloidin, magenta) and nuclei (DAPI, blue). Insets show in higher magnification the actin cables outlined in dashed rectangles. Arrowheads point to buckled actin filaments. Scale bars, 1 μm. **i,** Violin plot shows the angle (degrees) of deviation between actin cables and the vertical pore axis in parental control (*n* = 190) *and* SEPT9_i1 depleted (*n* = 185) U2OS cell regions (*n* = 3) confined in 8 μm wide pores. Statistical analysis was done with the Mann-Whitney U-test. ****, p < 0.0001. All violin plots show median (black line), first and third quartiles (colored lines). All images were acquired with super-resolution SoRa SDCM in pores with diameters of 8 μm, 5 μm and 3 μm wide pores 12 h, 24 h and 36 h, respectively, after seeding.

In addition to elevated zyxin, we observed that vertical actin cables were more bent in SEPT9-depleted cells (Fig. **2h-i**), possibly due to the compressive and tensile stresses of enhanced contractility as previously observed with reconstituted networks of non-sarcomeric actomyosin.^61^ We quantified actin cable orientation relative to the apicobasal axis of nuclear translocation by measuring the angle of deviation between actin cables and the vertical pore axis. Deviation angles were markedly higher in septin-depleted cells (Fig. **2i**). Collectively, these data indicate that actomyosin forces and mechanical strain are substantially elevated in the absence of SEPT9, suggesting that septins buffer the contractile forces of confined cell migration.

### Septins protect actin filaments from damage induced by myosin II hyperactivation

During confined 3D migration, a variety of forces - including frictional and hydrostatic - act on the nucleus and cytoskeleton, potentially damaging these structures through mechanical stress distinct from or additive to actomyosin-induced forces. We therefore sought to test whether septins buffer actomyosin forces using an orthogonal approach that relied on myosin II hyperactivation in the absence of additional mechanical constraints. We treated primary mouse embryonic fibroblasts (MEFs) from *Sept9*^-/-^ mice and littermate *Sept9^+/+^* controls^62^ (**Extended Data Fig. 4a,b**) with calyculin A, a potent inhibitor of protein phosphatase 1/2A that enhances phosphorylation of myosin regulatory light chain and consequently increases myosin II activity.^63^ We cultured cells on 2D collagen-coated substrates to exclude actomyosin-mediated effects from confinement-induced forces.

After treating MEFs with a low concentration of calyculin A (10 nM) for 15-30 minutes, we observed breaks in phalloidin-stained actin stress fibers (Fig. **3a**, arrowheads) proximal to peripheral focal adhesions, which are sites of higher mechanical tension. In *Sept9*^⁻/⁻^ MEFs, these breaks were more abundant (Fig. **3b**, arrowheads). Additionally, several actin stress fibers displayed a wavy appearance with multiple buckles (**Fig. 3b**, arrows). Quantification of the percentage of stress fibers with phalloidin breaks revealed that actin severing increased progressively during calyculin A treatment, with statistically significant increases between 0 and 30 minutes in both *Sept9^+/+^* and *Sept9*^-/-^ MEFs (Fig. **3c**). After 30 minutes, the percentage of severed actin filaments was markedly higher in *Sept9*^-/-^ than *Sept9^+/+^* MEFs (Fig. **3c**). Similarly, the percentage of buckled actin filaments was significantly elevated in *Sept9*^-/-^ compared to *Sept9*^⁺/⁺^ MEFs after both 15 and 30 minutes of calyculin A treatment (Fig. **3d**). Enhanced buckling without significant severing after 15 minutes was consistent with buckling preceding severing in early stages of contraction, and severing occurring on actin segments of high curvature.^61^ Notably, expression of human SEPT9_i1 - an isoform that crosslinks actin filaments and is required for stress fiber organization^41,44^ - in *Sept9*^⁻/⁻^ MEFs rescued the severing phenotype, reducing the percentage of severed actin filaments to control levels after 30 minutes of calyculin A treatment (Fig. **3e**).

**Fig. 3:**
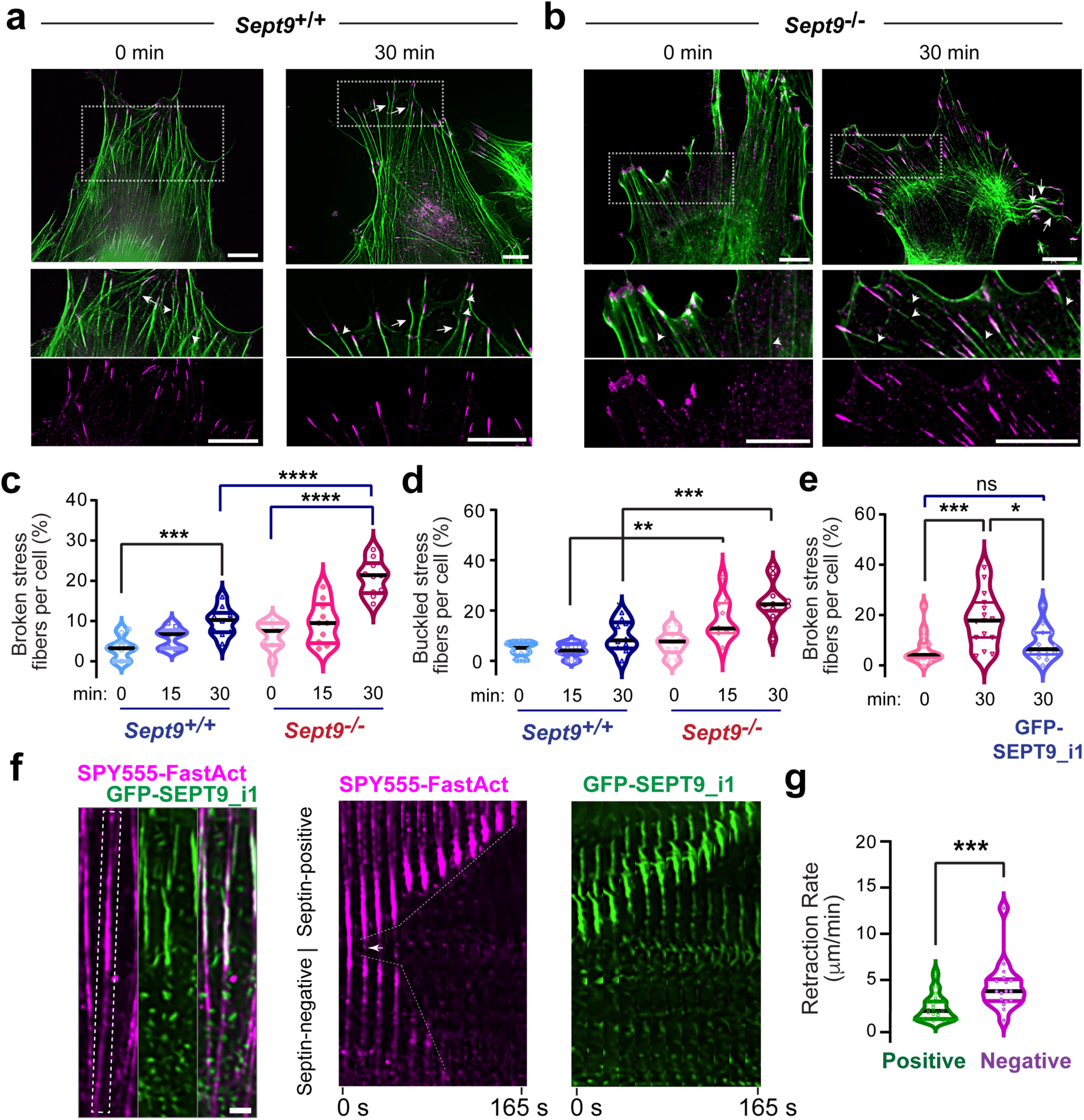
Septins relieve actin stress fibers from tensile stress induced by myosin II hyperactivation. **a-b**, Wide-field microscopy images show F-actin (phalloidin, green) and paxillin (magenta) in mouse embryonic fibroblasts (MEFs) from *Sept9*^+/+^ (a) and *Sept9*^-/-^ (b) mice, which were plated on collagen type I and treated with calyculin A (10 nM) for 30 minutes. Regions in dashed rectangles are shown in higher magnification below. Arrows and arrowheads point to breaks and buckles in actin stress fibers, respectively. Scale bars, 5 μm. **c,** Violin plots show percentage of actin stress fibers with breaks per cell after 0 (*n* = 230-289 stress fibers; 10 cells), 15 (*n* = 235-285 stress fibers; 10 cells) and 30 (*n* = 239-255 stress fibers; 10 cells) minute treatment of *Sept9*^+/+^ and *SEPT9*^-/-^MEFs with calyculin A (10 nM). Statistical comparisons between the indicated groups were done with Brown-Forsythe and Welch one-way ANOVA test with Dunnett T3 multiple comparisons test. ***, p<0.001; ****, p<0.0001. **d,** Violin plots show percentage of actin stress fibers with buckles per cell after 0 (*n* = 230-289 stress fibers; 10 cells), 15 (*n* = 235-285 stress fibers; 10 cells) and 30 (*n* = 239-255 stress fibers; 10 cells) minute treatment of *Sept9*^+/+^ and *Sept9*^-/-^ MEFs with calyculin A (10 nM). Statistical comparisons between the indicated groups were done with Brown-Forsythe and Welch one-way ANOVA test with Dunnett T3 multiple comparisons test. ***, p<0.001; ****, p<0.0001. **e,** Violin plots show percentage of stress fibers with breaks in untreated *Sept9*^-/-^ MEFs (0 min; *n* = 467; 15 cells), and in *Sept9*^-/-^ MEFs with (*n* = 425; 15 cells) or without GFP-SEPT9_i1 (*n* = 376; 15 cells) after 30 minutes of calyculin A (10 nM) treatment. Statistical comparisons between the indicated pair groups was done with the Mann-Whitney U-test. *, p<0.05; ***, p<0.001; ns, not significant. **f,** Spinning disk confocal microscopy image (left) and still frames (right) show a subcellular region of calyculin A (10 nM) treated *SEPT9*^-/-^ MEF cells, which expressed GFP-SEPT9_i1 (green) and labeled with SPY555-FastAct (actin stress fibers, magenta). Dashed rectangle outlines the actin fiber shown in higher magnification in still frames. Arrow points to point of severing between the GFP-SEPT9_i1-coated and uncoated region of the stress fiber. Dashed lines mark the retraction trajectories of the GFP-SEPT9_i1-free and -coated actin segments. **g,** Violin plots show the retraction rates (μm/min) for GFP-SEPT9_i1 free (*n* = 29) and coated actin stress fibers (*n* = 17). Statistical comparison was done with the Mann-Whitney U-test. ***, p<0.001. All violin plots show median (black line), first and third quartiles (colored lines).

To further examine whether SEPT9_i1 reduces mechanical tension on actin stress fibers, we sought to image and analyze actin filament severing in real time in living cells. We posited that if SEPT9_i1 buffers tension on actin filaments, severing would occur preferentially on SEPT9_i1-free actin stress fibers. We tested this prediction by transfecting GFP-SEPT9_i1 into *Sept9^-/-^* MEFs to generate a heterogeneous population of SEPT9_i1-coated and SEPT9_i1-free actin stress fibers, which were labeled with the vital dye SPY555-FastAct. Time-lapse imaging after calyculin A treatment revealed that 84% of severing events (*n* = 45) occurred on actin stress fibers segments lacking GFP-SEPT9_i1. Strikingly, on actin stress fibers that were partially coated with GFP-SEPT9_i1, the decorated segments retracted slower than the uncoated segments (Fig. **3f,g****; Supplementary Video 6**). The differential retraction velocity indicates that SEPT9_i1-coated segments are under lower tension. Collectively, these results suggest that SEPT9_i1 reduces tension on actin filaments, protecting them from severing under hypercontractile conditions.

### Septins mitigate mechanical stress on the nuclear envelope from actomyosin contractility

We reasoned that buffering of actomyosin tension by septins may extend to the nucleus, lessening the mechanical stress imposed on the nucleus during confined migration. To test if septins impact the forces transduced to the nuclear membrane by actomyosin contractility, we imaged the nuclear envelope during myosin II hyperactivation by calyculin A in *Sept9^+/+^* and *Sept9*^-/-^ MEFs cultured on 2D collagen. In control *Sept9^+/+^* MEFs, time-lapse imaging of LAP2β-AcGFP1 showed a gradual decrease in the nuclear area (Fig. **4a**). After 15 minutes of calyculin A treatment, regions of the circular nuclear envelope began to flatten (Fig. **4a**, white arrow) and crinkle forming small dimples (Fig. **4a**, white arrowheads). In contrast, myosin II hyper-activation had an earlier and more dramatic effect in *Sept9*^-/-^MEFs. The nuclear envelope began wrinkling after 10 minutes, and exhibited deep wrinkles and zigzagged indentations assuming a jagged morphology after an additional 5-10 minutes (Fig. **4a,b**).

**Fig. 4:**
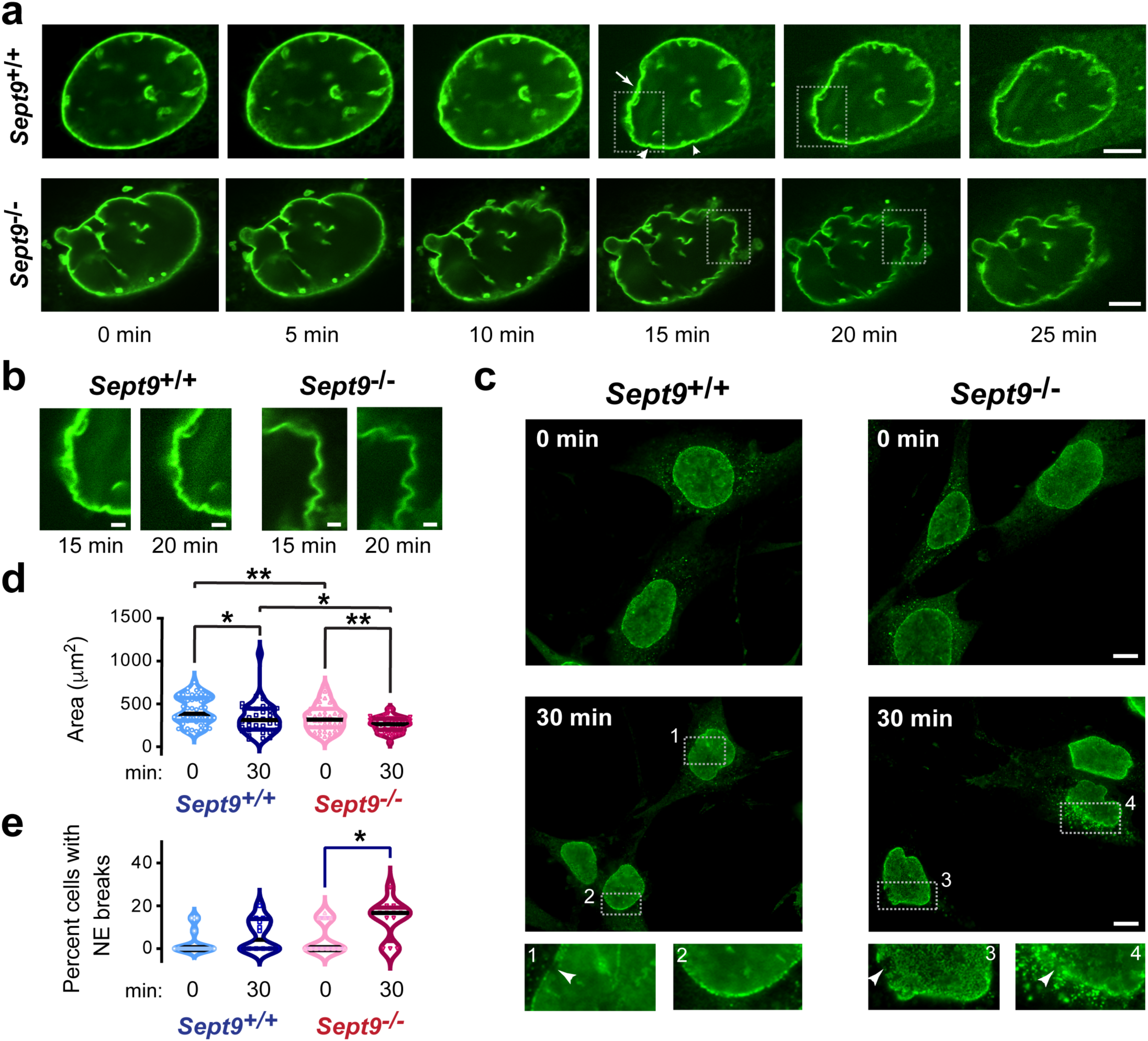
Septins protect the nuclear envelope from the mechanical stress of actomyosin contraction. **a**, Still frames of SDCM images of live Sept9+/+ and SEPT9-/- MEFs expressing LAP2β-AcGFP1 (green) and treated with calyculin A (10 nM) which was administered 5 minutes prior to start of time-lapse imaging. Arrow points to latten region of the circular nuclear envelope. Arrowheads point to nuclear envelope dimples. Scale bars, 5 μm. **b,** SDCM images show in higher magnification regions of the nuclear envelope outlined by the dashed lines in still frames of the time series above (a). Scale bars, 1 μm. **c,** Wide field microscopy images of *Sept9*^+/+^ and *Sept9^-/-^* immunostained for nucleoporins after 0 and 30 min of treatment with calyculin A (10 nM). Numbered regions outlined by dashed lines are shown in higher magnification. Arrows point to sites of nuclear envelope rupture. Scale bars, 10 μm. **d,** Violin plots show nuclear surface areas in *Sept9*^+/+^ MEFs after 0 minutes (*n* = 55 cells) and 30 minutes (*n* = 38 cells) of calyculin A treatment, and in *SEPT9*^-/-^ MEFs after 0 minutes (*n* = 50 cells) and 30 minutes (*n* = 50 cells) of calyculin A treatment. Statistical comparisons between the indicated groups were done with Brown-Forsythe and Welch ANOVA test (p<0.0001) with Dunnette’s T3 multiple comparisons test. *, p<0.05; **, p<0.01. **e,** Violin plots show percentage of cells with ruptured nuclear envelopes in *Sept9*^+/+^ MEFs after 0 minutes (*n* = 55 cells) and 30 minutes (*n* = 38 cells) of calyculin A treatment, and in *SEPT9*^-/-^ MEFs after 0 minutes (*n* = 50 cells) and 30 minutes (*n* = 50 cells) of calyculin A treatment. Statistical comparisons between the indicated groups were done with a non-parametric Kruskal-Wallis one-way ANOVA with Dunn’s multiple comparisons test. *, p<0.05. All violin plots show median (black line), first and third quartiles (colored lines).

To further assess and quantify the impact of actomyosin hypercontractility and SEPT9 depletion on the nuclear envelope, we immunostained MEFs for nucleoporins (Fig. **4c**). In agreement with real time observations of the LAP2β-AcGFP1-labeled nuclear membrane, nuclear envelopes were less circular in *Sept9^-/-^* MEFs, possessing jagged and polygonal shapes after 30 minutes of calyculin treatment (Fig. **4c**). Several nuclei exhibited herniation and discontinuous nuclear rims, indicative of rupture (Fig. **4c**, arrowheads). Cytoplasmic nucleoporin foci, likely remnants of nuclear pore complexes released after rupture, were also seen at nuclear envelope gaps (Fig. **4c**, inset 4 arrowheads). Quantification of nuclear areas in untreated *Sept9^+/+^* versus *Sept9^-/-^* MEFs showed a reduction in *Sept9^⁻/⁻^* MEFs (Fig. **4d**). After 30 minutes of calyculin A treatment, nuclear areas decreased in both *Sept9^+/+^* and *Sept9^-/-^* MEFs, and were lower in *Sept9^-/-^* compared to *Sept9^+/+^* (Fig. **4d**). Analysis of the percentage of cells with ruptured nuclear envelopes revealed a statistically significant increase in calyculin A-treated *Sept9^-/-^* MEFs, while *Sept9^+/+^* were unaffected (Fig. **4e**). Thus, loss of SEPT9 renders the nuclear envelope vulnerable to rupture, as unbuffered actomyosin forces exceed its mechanical tolerance.

### SEPT9_i1 protects the nucleus from genotoxic stress during confined migration, conferring survival advantage to breast cancer cells

The SEPT9 gene is amplified in murine models of breast cancer and human breast cancer cell lines.^47,64,65^ Importantly, the SEPT9_i1 isoform enhances cell migration^41,47–50^ and induces the lung metastasis of mammary cells in a murine xenograft model.^48^ Given that SEPT9_i1 expression alone was sufficient to reduce actomyosin tension in a SEPT9 pan-isoform knockout background (*Sept9^-/-^*MEFs; Fig. **3e**), we asked whether SEPT9_i1 amplification affects the mechanically induced genotoxic stress that cancer cells experience during confined migration.^20,21,66^

We first examined whether the highly invasive human breast cancer cell line MDA-MB-231,^67^ which harbors multiple SEPT9 gene copies,^47^ contains a perinuclear septoactomyosin cable arrangement similar to U2OS cells during confined migration. MDA-MB-231 cells possessed actin cables that surrounded the confined nucleus within narrow 3 μm-wide pores, aligning with SEPT9 along the apicobasal axis of the pore (**Extended Data, Fig. 5a, b; arrowheads; Supplementary Video 7**). Indicative of contractile activity, pMLC2 localized to several actin cables along with SEPT9 (**Extended Data, Fig. 5c-d**; arrowheads). Interestingly, the length of actin cables increased in pores with decreased diameter (**Extended Data, Fig. 5e,f**), while the overall number of actin cables per cell did not change despite the reduced volume of the confined cytoplasm (12-13 ± 4-6 cables; *n* = 9-10 cells) – indicating that more and longer actin cables form in narrower pores, potentially an adaptive response to the higher forces necessary to overcome increasing physical constraints.

We asked whether MDA-MB-231 cells are prone to increasing DNA damage after migration through pores of decreasing diameters, which is predicted to scale with increasing forces, and whether SEPT9_i1 expression affects the damage induced by mechanical stress. Using CRISPR, we generated stable SEPT9_i1-depleted MDA-MB-231 cells (**Extended Data, Fig. 5g-h**). We assessed DNA lesions in cells that had fully emerged onto the basal membrane surface after 36 h of EGF-guided chemotactic migration. Of note, the basal side was coated with the same ECM type that filled the pores and coated the apical side. Immunostaining for γH2AX, a marker of DNA double-strand breaks (DSBs), revealed a marked increase in the percentage of control cells with large γH2AX foci among those that emerged from 3 μm pores compared to 5 μm and 8 μm wide pores (Fig. **5a-c**). Strikingly, DNA damage increased in SEPT9_i1-depleted MDA-MB-231 compared to controls after confined migration through 8 μm, 5 μm, or 3 μm wide pores (Figure **5c**).

**Fig. 5:**
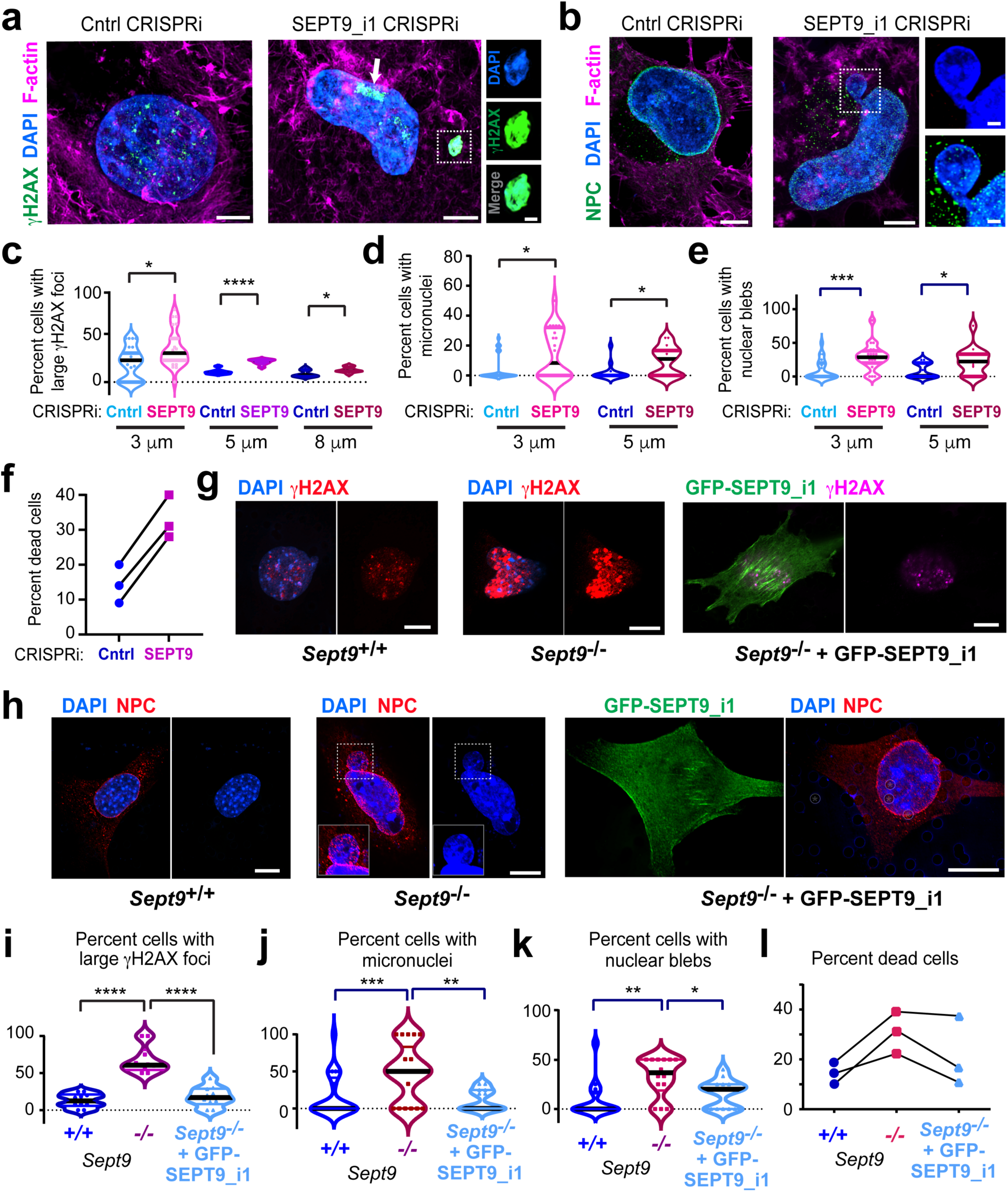
SEPT9_i1 protects nuclei from mechanical damage during confined migration, suppressing DNA damage, nuclear defects associated with genomic instability, and cell death. **a**, SoRa SDCM images of nuclei (DAPI, blue), γH2AX (green) and actin (phalloidin, magenta) in parental control and SEPT9_i1-depleted MDA-MB-231 cells, taken from the basal side of polycarbonate membranes with 3 μm wide pores after 36 h of apicobasal transmigration. Arrow points to a large nuclear region with γH2AX accumulation. Inset shows a γH2AX-enriched micronucleus in higher magnification. Scale bars, 5 μm (1 μm in inset). **b,** SDCM images of nuclei (DAPI, blue), nucleoporins (green) and actin (phalloidin, magenta) in parental control and SEPT9_i1-depleted MDA-MB-231 cells, taken from the basal side of polycarbonate membranes with 3 μm wide pores after 36 h of apicobasal transmigration. Inset shows in higher magnification a region of nuclear blebbing, where nuclear DNA protrudes from a site of nuclear envelope rupture. Scale bars, 5 μm (1 μm in inset). **c,** Violin plots show percentage of parental control and SEPT9_i1-depleted MDA-MB-231 cells with large γH2AX foci on basal membrane regions 3 μm (*n* = 26 regions; 83-96 cells), 5 μm (*n* = 9 regions; 74-89 cells) and 8 μm (*n* = 9 regions; 80-106 cells) wide pores. Statistical comparisons between the indicated pair groups were done with the Mann Whitney U-test. *, p<0.05; ****, p<0.0091. **d,** Violin plots show percentage of parental control and SEPT9_i1-depleted MDA-MB-231 cells with micronuclei on basal membrane regions with 3 μm (*n* = 22 fields; 88-95 cells) and 5 μm (*n* = 15 fields; 80-85 cells). Statistical comparisons between the indicated pair groups was done with the Mann Whitney U-test. *, p<0.05. **e,** Violin plots show percentage of parental control and SEPT9_i1-depleted MDA-MB-231 cells with nuclear blebs on basal membrane regions with 3 μm (*n* = 22 fields; 88-95 cells) and 5 μm (*n* = 15 fields, *n* = 80-85 cells) pores. Statistical comparisons between the indicated pair groups was done with the Mann Whitney U-test. *, p<0.05; ***, p<0.001. **f,** Percentage of Trypan blue-positive cells after trypsinization of parental control (*n* = 5230, 3660, 2620) and SEPT9_i1-depleted MDA-MB-231 cells (*n* = 4190, 3140, 2090) from the basal surfaces of Transwell polycarbonate membranes with 3 μm wide pores, following 36 h of apicobasal transmigration. Results from three independent experiments are shown. **g,** SDCM images of nuclei (DAPI, blue), γH2AX (red) and GFP-SEPT9_i1 in *Sept9*^+/+^ and/or *Sept9*^-/-^ MEFs, taken from the basal side of polycarbonate membranes with 3 μm wide pores after 36 h of apicobasal transmigration. Scale bars, 10 μm. **h,** SDCM images of nuclei (DAPI, blue), nucleoporins (red) and actin (phalloidin, magenta) in *Sept9*^+/+^ and/or *Sept9*^-/-^MEFs cells, taken from the basal side of polycarbonate membranes with 3 μm wide pores after 36 h of apicobasal transmigration. Scale bars, 10 μm. **i,** Violin plots show percentage of cells with large γH2AX foci after imaging basal regions of polycarbonate membranes with 3 μm wide pores containing *Sept9*^+/+^ MEFs (*n* = 22 regions; 48 cells) and *Sept9*^-/-^ MEFs with (*n* = 22 regions; 35 cells) and without GFP-SEPT9_i1 (*n* = 12 regions; 63 cells). Statistical comparisons between the indicated groups were done with Brown-Forsythe and Welch ANOVA test with Dunnett T3 multiple comparisons test. ****, p<0.0001. **j,** Violin plots show percentage of cells with micronuclei after imaging basal regions of polycarbonate membranes with 3 μm wide pores containing *Sept9*^+/+^ MEFs (*n* = 22 regions; 48 cells) and *Sept9*^-/-^ MEFs with (*n* = 22 regions; 35 cells) and without GFP-SEPT9_i1 (*n* = 12 regions; 63 cells). Statistical comparisons between the indicated groups were done with Brown-Forsythe and Welch ANOVA test with Dunnett T3 multiple comparisons test. **, p<0.01; ***, p<0.001. **k,** Violin plots show percentage of cells with nuclear blebs after imaging basal regions of polycarbonate membranes with 3 μm wide pores containing *Sept9*^+/+^ MEFs (*n* = 14 regions; 51 cells) and *Sept9*^-/-^ MEFs with (*n* = 14 regions; 45 cells) and without GFP-SEPT9_i1 (*n* = 14 regions; 51 cells). Statistical comparisons between the indicated groups were done with Brown-Forsythe and Welch ANOVA test with Dunnett T3 multiple comparisons test. **, p<0.01; *, p<0.05. All violin plots show median (black line), first and third quartiles (colored lines). **l,** Percentage of Trypan blue-positive cells after trypsinization of *Sept9*^+/+^ (*n =* 4115, 2939, 4700), *Sept9*^-/-^ (*n =* 6760, 5580, 2645) and *Sept9*^-/-^ MEFs transfected with GFP-SEPT9_i1 (*n =* 2175, 1764, 7055) for 24 h prior to confined migration for 36 h through 3 μm wide pores. Results from three independent experiments are shown. Note that transfection efficiency was ∼40%, and cell death was measured in the entire population of *Sept9*^-/-^ MEFs and not strictly in the cells expressing GFP-SEPT9_i1. All violin plots show median (black line), first and third quartiles (colored lines).

We next analyzed cells for micronuclei and nuclear blebs, biomarkers of genomic instability resulting from excessive DNA damage and nuclear rupture.^23^ We stained cells for nucleoporins as well as DAPI to visualize DNA, quantifying micronuclei as cytoplasmic islands of DNA lacking nucleoporins (Fig. **5a**, inset) and nuclear blebs as nuclei with DNA bulging out through discontinuities in the nuclear envelope (Fig. **5b**, inset). We focused this analysis on cells that emerged from narrower 3-5 μm wide pores, which induce higher mechanical stress. SEPT9_i1-depleted cells showed a statistically significant increase in both micronuclei and nuclear blebs after migration through 3 μm and 5 μm pores (Fig. **5d,e**). Notably, the percentage of cells with nuclear blebs was markedly higher than those with micronuclei, with nearly 25% of SEPT9_i1-depleted cells containing nuclear blebs (Fig. **5e**). As micronuclei typically form after mitosis, the higher prevalence of nuclear blebs may reflect a greater proportion of cells that had not yet undergone cell division after nuclear rupture.

Because SEPT9_i1-depleted cells exhibit increased DSBs, micronuclei, and nuclear blebs - all hallmarks of mechanical genotoxic damage - we asked whether cells that emerge from confined migration are less viable. To assay cell death, we harvested cells adhered to the basal surface of the polymeric membranes following confined migration through 3 μm wide pores and labeled them with Trypan Blue to distinguish live from dead cells. We found that 25-40% of SEPT9_i1-depleted cells were dead, representing a 2-3-fold increase compared to control MDA-MB-231 cells (Fig. **5f**). Similar results were obtained independently using a different SEPT9_i1-depleted MDA-MB-231 clone, which also exhibited increased DNA damage, micronuclei, nuclear blebs and cell death after confined migration (**Extended Data, Fig. 5i-l**).

We next asked whether the oncogenic SEPT9_i1 isoform alone can limit the genotoxic mechanical damage which is caused by confined migration. Because SEPT9_i1 expression rescued actin stress fiber damage in *Sept9*^⁻/⁻^ MEFs under hypercontractile conditions induced by myosin II hyperactivation, restoring actin stress fiber integrity to control levels (Fig. **3e**), we tested whether SEPT9_i1 expression can suppress genotoxic damage and cell death during confined migration in MEF cells. Following confined migration through 3 μm pores, the percentage of *Sept9^-/-^* MEFs with DDSBs, micronuclei, and nuclear blebs was substantially elevated compared to *Sept9^+/+^* MEFs (Fig. **5g,h**). Approximately 50% of *Sept9*^⁻/⁻^MEFs exhibited DNA damage and micronuclei (Fig. **5i-j**), while ∼30% contained nuclear blebs (Fig. **5k**). Additionally, cell death increased (Fig. **5l**). Expression of GFP-SEPT9_i1 in *Sept9^-/-^* MEFs rescued DNA damage and micronuclei, restoring the percentage of affected cells to *Sept9^+/+^* levels (Fig. **5i-j**), and also reduced nuclear blebs and cell death (Fig. **5k,l**). Taken together, these results show that SEPT9_i1 protects cells from the consequences of confined migration - DNA damage, genomic instability, and cell death – providing a survival advantage. Thus, the function of SEPT9_i1 in buffering actomyosin forces extends to protecting the nuclear and genomic integrity.

### *SEPT9* amplification associates with tolerance of genomic instability in aggressive tumors and low patient survival

Metastatic cancers are characterized by elevated genomic instability, which promotes tumor evolution but also imposes constrains on cancer cell viability.^68–70^ Excessive genomic alterations compromise cell fitness and can trigger apoptosis and immunogenic cell death.^68,69^ During metastatic dissemination, which involves confined migration, tumor cells are subjected to recurrent nuclear and DNA damage during confined migration, increasing the likelihood that highly altered genomes exceed tolerable limits of genomic instability.^71^ This points to cellular mechanisms that permit continued tumor survival by limiting catastrophic genomic damage as evidenced by the low survival of tumors with excessive genomic alterations, the persistence of early (truncal) oncogenic mutations throughout cancer progression, and the presence of islands of relative genomic stability in metastatic tumor subclones.^69,72,73^

Our results suggest that SEPT9_i1 protects cancer cells from the genotoxic mechanical stress of confined migration. *SEPT9* amplification may therefore enable cancer cells to tolerate substantial genomic instability within a permissive threshold, potentially without preventing the accumulation of smaller alterations, which might be beneficial for tumor progression. To assess whether this model is reflected in clinical outcomes, we examined genomic and survival data from cancer patients.

Systematic analysis of *SEPT9* gene copy number alterations in large clinicgenomic cohorts are lacking, and the relationship between SEPT9, genomic alterations and patient outcomes has not been explored. We analyzed overall survival data in the Cancer Genome Atlas (TCGA), comparing pan-cancer patients with *SEPT9* gene copy number gain (3-4 copies; +1) or amplification (>5 copies; +2) to copy neutral tumors. Strikingly, patients with *SEPT9* gain or amplification exhibited reduced overall survival (Fig. **6a**).

**Fig. 6:**
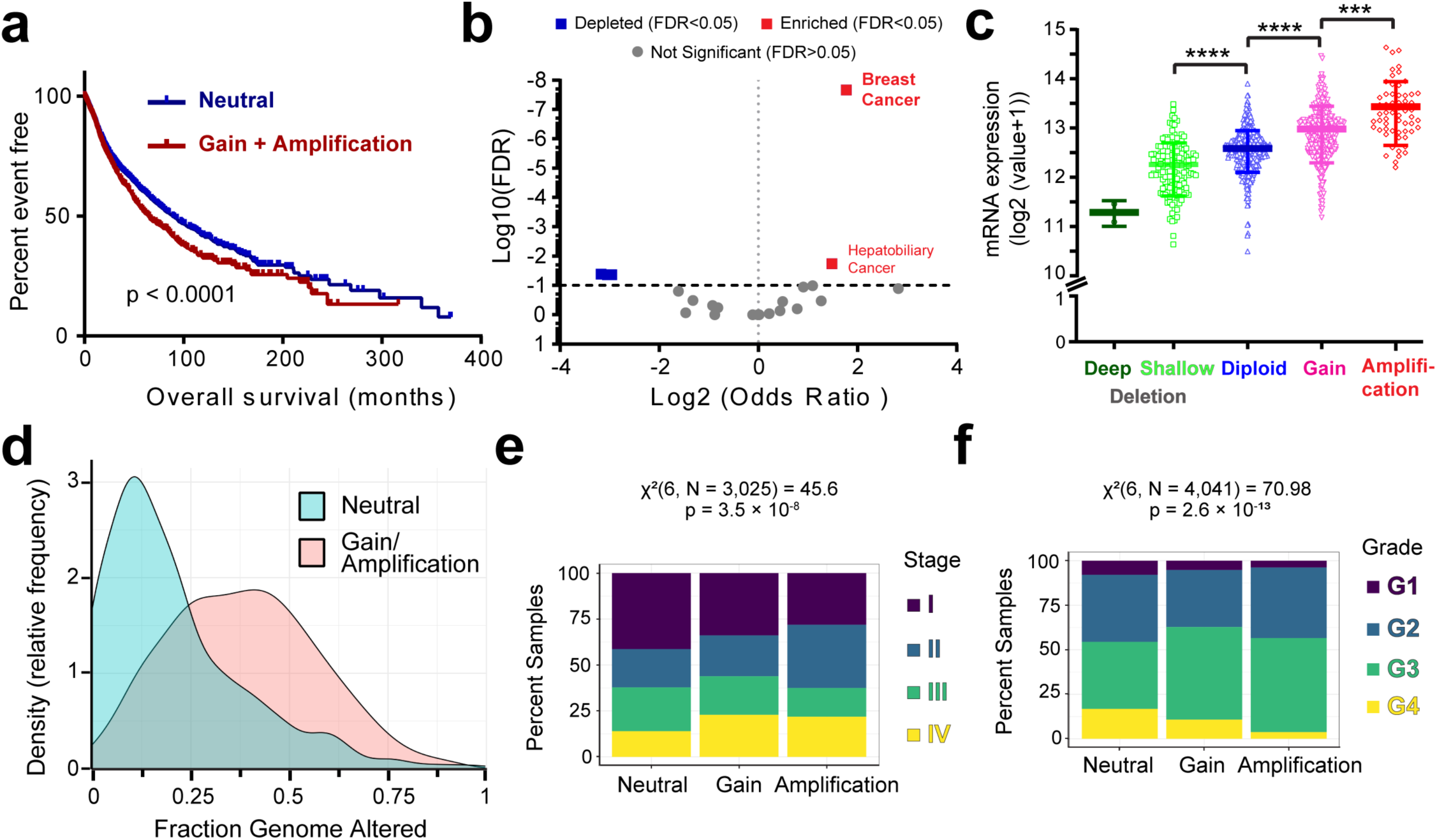
*SEPT9* amplification associates with suppression of excessive genomic instability and lower patient survival. **a**, Kaplan-Maier plot of survival in a pan-cancer TCGA cohort stratified by *SEPT9* copy number status. Patients with copy-neutral (diploid) *SEPT9* tumors (blue; *n* = 6,912) are compared with patients whose tumors harbor SEPT9 copy-number gain or amplification (red; *n* = 2,873). Survival curves were compared using the log rank test (p < 0.0001; *n* = 3,135 events). **b,** Volcano plot showing log2 odds ratios for *SEPT9* amplification (positive values) and deletion (negative values) across tumor types, plotted against statistical significance (-log10 FDR). Tumor types with FDR < 0.05 are highlighted. Breast cancer shows the strongest enrichment for *SEPT9* amplification, while more modest enrichment is observed in hepatobiliary cancer. Several tumor types including (blue squares) glioblastoma, head and neck cancer, thyroid cancer, and renal clear cell carcinoma, exhibit significant depletion of *SEPT*9 copy number. Data were derived from TCGA. **c,** SEPT9 mRNA expression (log2 (value+1)) is shown for breast tumors stratified by *SEPT9* GISTIC copy-number state: deep (*n* = 2) and shallow deletion (*n* = 153), diploid (*n* = 447), gain (*n* = 299), and amplification (*n* = 59). Statistical analysis was assessed using Kruskal-Wallis test (p < 0.0001), followed by Dunn’s multiple comparisons test (****, p < 0.0001; ***, p < 0.001). **d,** Kernel density plot showing the relative frequency of fractional genome altered (FGA) values in breast tumors with copy-neutral *SEPT9* (cyan; *n* = 517) and tumors with *SEPT9* copy-number gain or amplification (pink; *n* = 403). **e,** Stacked bar plot showing the distribution of simplified tumor stages (I–IV) across *SEPT9* copy-number states (neutral, gain, amplified) in primary tumors from TCGA. Proportions are calculated within each *SEPT9* copy-number category. *SEPT9* gain and amplification are progressively enriched among higher-stage tumors. Statistical significance was assessed using a chi-square test (χ²(6, N = 3,025) = 45.64, p = 3.5 × 10⁻⁸). **f,** Stacked bar plot depicting the distribution of histologic tumor grades (G1-G4) stratified by *SEPT9* copy-number status across primary tumors. Only samples with interpretable grade annotations (G1–G4) were included. *SEPT9* gain and amplification are associated with increased proportions of higher-grade tumors. Statistical significance was evaluated using a chi-square test (χ²(6, N = 4,041) = 70.98, p = 2.6 × 10⁻¹³). G1, well differentiated (low grade); G2, moderately differentiated (intermediate grade); G3, poorly differentiated (high grade); G4, undifferentiated (high grade).

To assess tumor-type specificity of *SEPT9* copy-number alterations, we next examined *SEPT9* amplification frequency across all TCGA primary tumors with available copy-number data (*n* = 11,245). *SEPT9* amplification (>5 copies) was most frequent in invasive breast, renal, and hepatic cancers, occurring in approximately 25% of cases within each cohort (**Extended Data, Fig. 6a**). To formally evaluate tumor-type enrichment of high-level *SEPT9* amplification, we performed enrichment analyses restricted to amplification-only events in tumor types with sufficient sample size (N ≥ 20). Breast cancer emerged as the most significantly enriched tumor type for *SEPT9* amplification (odds ratio = 3.42, BH–FDR = 2.16 × 10⁻⁸), with more modest enrichment observed in hepatobiliary cancers (odds ratio = 2.80, BH–FDR = 0.018) (Fig. **6b**).

Because gene copy-number alterations do not necessarily imply functional relevance, we next examined *SEPT9* mRNA expression in breast cancer tumors stratified by *SEPT9* gene copy-number status. Mean *SEPT9* mRNA expression increased in a stepwise manner with increasing *SEPT9* gene copy number, including copy-number gain and amplification (Fig. **6c**). This concordance between gene copy-number increase and transcriptional upregulation is consistent with dosage-dependent regulation and mirrors patterns observed for established oncogenes in breast cancer, such as *ERBB2* (*HER2)*.^74,75^

To determine whether *SEPT9* gene copy gain and amplification simply reflect elevated chromosomal instability, we examined the distribution of fractional genome altered (FGA) across tumors stratified by *SEPT9* copy-number status in breast cancers. The relative frequency of *SEPT9* gain and amplification varied across the full spectrum of FGA values, exhibiting a non-monotonic distribution that peaked at intermediate levels of chromosomal instability (FGA ∼0.25-0.5) and declined at higher FGA values (0.5-0.7) (Fig. **6d**). Notably, the FGA range over which *SEPT9* gain and amplification were most frequent overlaps with levels of chromosomal instability commonly observed in aggressive breast cancer subtypes with higher recurrence rates and adverse clinical outcomes, including luminal B and basal/triple-negative tumors (**Extended Data Fig. 6b**). A pan-cancer analysis also revealed a strong relationship between *SEPT9* copy-number state (neutral, gain, amplification) and tumor stage and grade, with *SEPT9* gain and amplification progressively enriched in tumors with higher stage and histologic grades (Fig. **6e,f**). These data indicate that *SEPT9* gene amplification associates with aggressive tumors with intermediate-to-high levels of genome alteration, which have a survival advantage over tumors with low or excessive levels of chromosomal instability.

If *SEPT9* amplification enables tumor cells to tolerate genomic instability, a key question is whether this tolerance involves selective constrains that limit the accumulation of larger scale genomic changes while allowing lower scale alterations. We examined this by analyzing the relationship between *SEPT9* copy-number status and tumor mutation burden (TMB), a measure of smaller scale genomic alterations from single nucleotide and small insertion-deletion events.

In contrast to FGA, *SEPT9* gain or amplification was not associated with reduced mutation burden. In tumor types enriched for *SEPT9* copy-number alterations, tumors harboring *SEPT9* gain or amplification exhibited comparable or modestly higher log-transformed TMB compared to copy-neutral tumors, whereas no significant difference was observed in pan-cancer analyses. In addition, *SEPT9* gain or amplification was not associated with a reduced probability of hypermutation, which was defined as tumors in the upper tail of the TMB distribution within each cancer type (top 20%). In tumor types enriched for *SEPT9* copy-number alterations, hypermutation was observed at comparable or higher frequencies in *SEPT9*-altered tumors compared to copy-neutral tumors (odds ratio = 1.65, FDR < 1 × 10⁻⁴), whereas no enrichment or depletion was observed in pan-cancer analyses.

Collectively, these results indicate that *SEPT9* gain and amplification are not simply a byproduct of extreme genome-wide chromosomal instability, and do not appear to associate with lower mutational burden. Instead, *SEPT9* amplification appears to enable tumor cells to tolerate substantial chromosomal instability without crossing lethal thresholds of genomic damage, and thereby may provide a selective advantage for aggressive tumors in certain metastatic niches. In this model (Fig. **7**), *SEPT9* protects the nucleus from catastrophic damage during confined migration while permitting continued accumulation of smaller-scale genetic alterations that can contribute to tumor heterogeneity, clonal evolution, and reduced clinical outcome.^24^

**Fig. 7:**
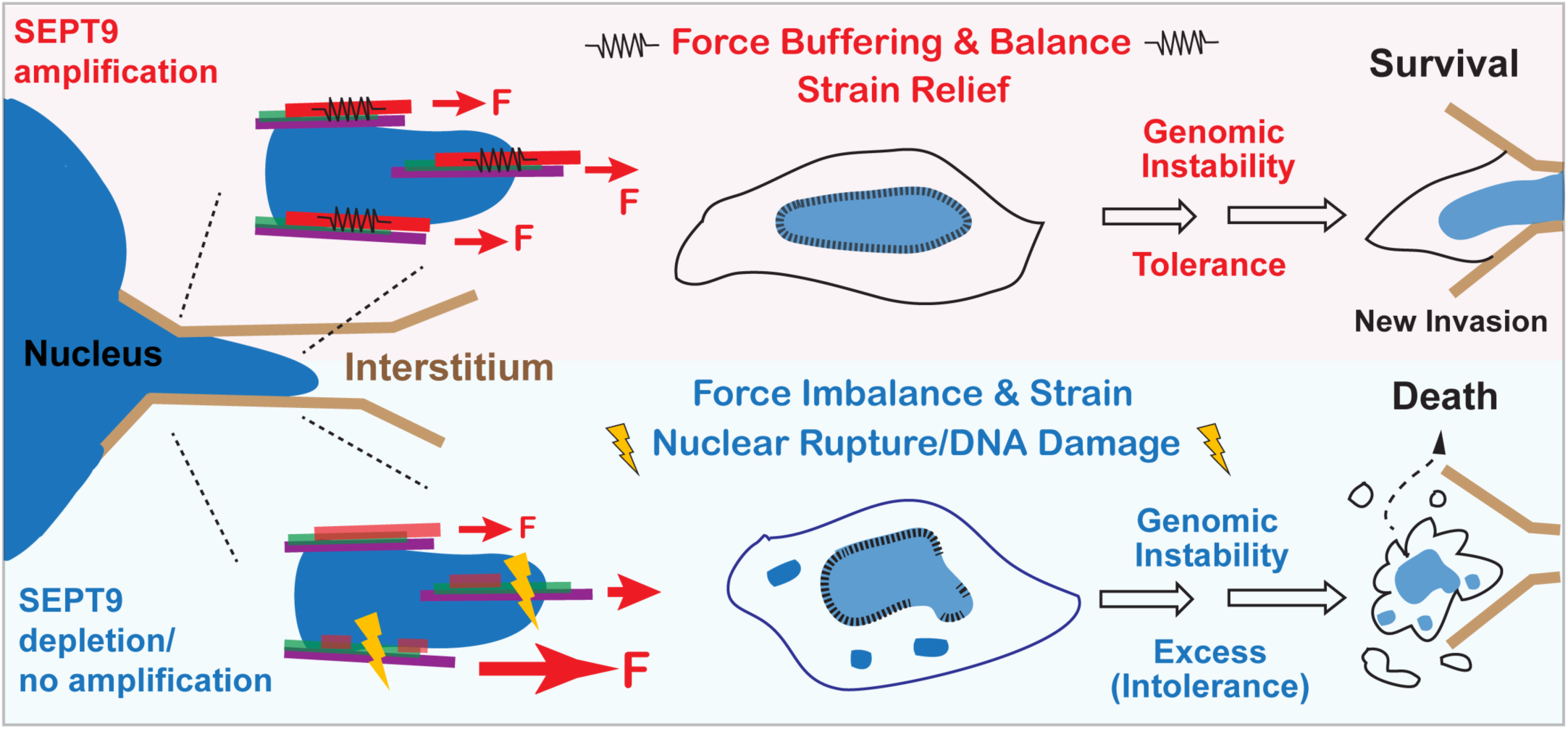
Working model: *SEPT9* amplification buffers mechanical forces during confined migration to maintain a permissive window of genomic instability. The schematic illustrates the proposed role of *SEPT9* amplification in buffering actomyosin-generated forces during confined migration to maintain genomic instability within tolerable limits. We posit that through this mechanism, *SEPT9* amplification enables metastatic tumor cells to navigate a narrow survival window, avoiding lethal genomic catastrophe, which ultimately promotes metastatic colonization and diminishes patient survival. In tumor cells with lower levels of SEPT9 expression, the nucleus is more prone to mechanical damage during confined migration, leading to large scale genomic alterations that surpass a threshold of cell tolerance and result in cell death.

## Discussion

Septins comprise a distinct cytoskeletal network - often referred to as the fourth component of the cytoskeleton alongside actin, microtubules, and intermediate filaments.^35^ While septins have been reported to influence plasma membrane rigidity and the nucleocytoplasmic transport of mechanosensitive transcription factors,^76,77^ they have been largely overlooked as a mechanical element of the cell. Here, we reveal a previously unknown septin function in buffering actomyosin forces, and thereby protecting both actin cytoskeleton and the nucleus from mechanical stress and damage.

Septins are neither evolutionarily nor structurally related to intermediate filaments, which are flexible and stretchable structures that dissipate mechanical stress through deformation and reorganization.^33,34^ This raises the question of how septins can buffer mechanical tension. Intriguingly, yeast septin filaments have persistence lengths of ∼1-2 μm, approaching those of the pliable intermediate filaments (0.2-1 μm).^78,79^ Septins’ persistence lengths are approximately one and three orders of magnitude lower than those of actin filaments (∼10-20 μm) and microtubules (∼5,000 μm), respectively.^80^ Importantly, septin filaments possess both longitudinal and lateral plasticity. The oligomeric interfaces of human septins can compact and expand in a nucleotide-dependent manner (GTP/GDP),^81^ and can hinge up to 30 degrees between the terminal subunits of their protomeric oligomers.^82^ Septin filaments also interact laterally through C-terminal coiled coils of variable length that extend sideways from the linear multimer.^83,84^ Resembling the side chains of bottlebrush polymers used in elastic biomaterials and electronics,^85^ these C-terminal extensions may enable septin filaments to expand, contract, and rearrange lateral contacts within filament bundles in response to intracellular forces, assembling into mechanically resilient networks.^83^ Overall, the structural plasticity of septin filaments makes them well-suited to absorb and dissipate mechanical strain.

Septins may absorb forces not only on their own as deformable semi-elastic polymers, but also as components of actomyosin networks. Actin binding and crosslinking proteins (e.g., anillin, filamin, fascin) modify the rheological properties and behavior of actin networks in response to mechanical strain.^86–89^ Depending on the strength of interaction with actin filaments, protein crosslinkers can dynamically rearrange in response to stress, impacting actin network mechanics. Septins crosslink actin filaments via SEPT9^41,42,44,45^ and therefore, may affect the response of actin networks to mechanical stress. Previous work showed that septin complexes crosslink actin filaments in vitro into curved and circular bundles,^90^ indicating that septins can reduce the rigidity of actin polymers, possibly by inducing conformational changes and flexible links that enable bending without breaking. Therefore, septin presence in actomyosin networks may reduce flexural rigidity, and thereby, minimize severing under higher mechanical loads. In retrospect, this function of septins may account for the disassembly and disorganization of highly contractile actin networks such as actin stress fibers and cytokinetic rings, which have been extensively reported in a diversity of cell types and organisms.^37^

During confined cell migration, the nucleus is subjected to contractile, tensile and hydrostatic forces that enable translocation through spaces smaller than its size.^91^ Migratory cells adapt different strategies for threading their nucleus through tight spaces, and they have mechanisms for repairing nuclear envelope rupture and DNA damage. However, mechanisms that cushion the nucleus from the mechanical impact of rising intracellular forces remain poorly understood. The intermediate filament protein vimentin, which is upregulated in invasive cancer cells undergoing epithelia-to-mesenchymal transition, forms a perinuclear cage that protects against nuclear deformation.^30^ The microtubule network also provides a cushion around the nucleus, bearing the load of contractile forces and cross-talking with the actomyosin cytoskeleton by feedback signaling.^31^ Vimentin, which associates with microtubules via plectin, and the microtubule-associated proteins CLASP reinforce the microtubule network to withstand higher intracellular forces.^92^ While vimentin can absorb contractile forces on its own or in association with microtubules, it is also conducive to tensile force transmission, promoting the traction forces of cell migration.^32,34^

Septins emerge as cytoskeletal polymers that protect the nucleus by relieving tensile actomyosin forces in a manner, which is distinct from vimentin. During confined 3D cell migration, we found that septins do not form a perinuclear basket like vimentin but instead co-align with actomyosin cables that surround the nucleus axially in the apicobasal direction of movement, and connect with focal adhesions. On 2D substrates, septin depletion increases actin stress fiber severing at sites proximal to focal adhesions in response to myosin II hyperactivation. Notably, severing occurs preferentially on septin-free actin stress fibers, which retract faster than septin decorated actin filaments, indicating higher tension. Taken together with an increase in nuclear envelope rupture events in septin depleted cells, these results suggest that septins buffer the actomyosin forces that move the nucleus forward during confined migration. Interestingly, SEPT9 levels on actin cables increase along with myosin regulatory light chain phosphorylation with increasing confinement (narrower pores) and higher tensile stress (Fig. **2f,g**). This might be a mechanical adaptive response for threading the nucleus forward by a controlled and sustained pulling, where forces are uniformly spread and balanced (Fig. **7**) - analogous to how the suspension cables of a hot air balloon distribute lifting forces evenly across the passenger basket. This type of nuclear movement may prevent forces from concentrating focally at sites of the nuclear envelope, which can cause nuclear rupture and DNA damage. Septoactin cables, however, surround the confined nucleus without making any ostensible direct end-on contacts, suggesting that septins may buffer force waves that sweep tangentially along the nuclear envelope (i.e., perpendicular to the nuclear radius).

Septins localize preferentially to membrane domains experiencing elevated contractile force and tension, such as the cleavage furrow and cell junctions,^93–96^ as well as regions where force distribution and buffering are critical - including T cell mid-zones that suppress protrusive activity^97^ and septoactomyosin cages that constrain the motility of pathogenic bacteria.^98^ The preferential association of septins with domains of micron-scale membrane curvature may serve a similar protective function by relieving tension from membrane stretching at the base and neck of membrane protrusions, or by counteracting turgor pressure at fungal protrusions such as the hyphae of *Aspergillus* and the appressorium of the rice blast fungus.^99,100^

In work submitted concurrently with this manuscript, Utgaard et al. discovered that septins accumulate on the cell cortex in response to local mechanical compression and signaling.^101^ Our combined results suggest that septins can absorb mechanical strain in the membrane, potentially acting as a relief valve for the cytoskeleton and membranes that is conserved across diverse cell types and biological systems. Future work will explore whether tension, and specific tension thresholds, are critical determinants of septin assembly at membrane domains and actomyosin filaments.

In cancer cells undergoing invasive migration, mechanical tension and stress impact survival not only by compromising nuclear and genomic integrity but also by signaling to apoptotic pathways, which can induce programmed cell death.^20,22,23,102^ While external microenvironments dictate the types of forces and mechanical stresses induced, how tumor cells mitigate and adapt to these pressures can determine pathogenic course and survival outcomes. Our results suggest that SEPT9_i1 enhances survival by reducing genotoxic mechanical stress. We posit that this function provides a selective advantage for tumor clones and/or subclones suited for invasion and/or growth in specific niches by preserving their highly altered genomes during dissemination through confined tissue spaces. Thus, *SEPT9* amplification may enable metastatic tumor cells to navigate a narrow survival window - buffering actomyosin-generated forces during confined migration to maintain genomic instability within tolerable limits while avoiding lethal genomic catastrophe. ^68,69^ Interestingly, recent work showed that in non-adherent cancer cells (melanomas), septins scaffold survival signaling pathways on the curvatures of the blebbing plasma membrane.^103^ Taken together with the findings of this and the companion manuscript by Utgaard et al,^101^ septins appear to adapt pro-survival functions on the actomyosin cytoskeleton and cell membrane in a morphodynamic and mechanoresponsive manner, which in turn depends on tumor microenvironment and cell-ECM interactions. Overall, septin amplification emerges as a critical component of the pro-survival toolbox of cancer, raising the value of septins as precision oncology biomarkers and therapeutic targets.

## Methods

### Cell culture

#### Maintenance

The mammalian cell lines U2OS (ATCC HTB-96) and MDA-MB-231 (ATCC HTB-26) were maintained in DMEM High Glucose medium (Sigma-Aldrich-07777). Primary mouse embryonic fibroblasts were derived from *Sept9*^-/-^mice and control *Sept9*^+/+^ littermates, and immortalized as previous described. They were gifted by and Dr. Ernst-Martin Füchtbauer (Aarhus University, Denmark).^62^ All media were supplemented with 10% fetal bovine serum (Atlas Biologicals FR-0500-A) and 1% PSK (penicillin-streptomycin-kanamycin solution). Cells were maintained in culture at 37 °C in a humidified incubator with 5% CO₂. Cultures were maintained at 70% confluence and sub-cultured at a 1:5–1:10 ratio every 4 days using 0.05% trypsin-EDTA (Gibco, 25300-054).

#### Plating

Prior to experiments, cells were seeded on either glass coverslips (22 × 22 mm, No. 1.5; VWR, 48366-227), which were coated with type I collagen (PureCol type I bovine collagen; Advanced Biomatrix, 5005), or transparent polyester (PET) membrane Transwell inserts (Corning) coated with Matrigel and laminin. Glass coverslips were coated with collagen at a concentration of 2.5 μg/ml for 5 minutes at room temperature, and subsequently were air-dried for 30 minutes and exposed to UV light for 30 minutes in a biosafety cabinet to ensure sterilization. The PET membranes of Transwell inserts were coated on both sides with 200 µm of ice-cold ∼10 mg/mL Matrigel (Corning, 356237) and 0.2 mg/ml laminin (Sigma, L2020) on both sides and incubated at 4°C with gently rocking for 30 minutes to ensure even deposition. Excess Matrigel was aspirated, and membranes were polymerized at 37 °C for 30 minutes in a cell culture incubator prior to cell seeding. Membranes were then hydrated on both sides with DMEM High Glucose medium supplemented with 10% FBS and 1% PSK.

For experiments performed on glass coverslips, cells were detached using 0.05% trypsin-EDTA for 5 minutes at 37 °C, neutralized with complete DMEM (DMEM High Glucose medium, 10% fetal bovine serum and 1% PSK), and collected by centrifugation at 1,500 rpm for 5 minutes. The cell pellet was resuspended in fresh complete DMEM, and viable cells were counted using an automated cell counter Countess 3 (Thermo Fisher Scientific) with 0.4% (w/v) trypan blue exclusion (Corning, 25-900-Cl). Cells were seeded at a density of 2 × 10⁵ cells per coverslip and allowed to adhere and recover for 12–24 hours before transfection. For experiments using 24 mm Transwell membrane inserts, cells were seeded at a density of 1 × 10⁶ cells per insert.

For live cell imaging experiments, cells were seeded onto collagen-coated 35 mm glass No 1.5 bottom dishes (MatTek, P35G-1.5-20-C) and switched to phenol red-free HyClone DMEM/High Modified media (SH30284.01) prior to transferring to the microscope’s stagetop incubator. Live imaging of cells during confined migration through the pores of Transwell membranes was performed by cutting a rectangle strip from the Transwell’s membrane insert and immobilizing it on the glass portion of a MatTek 35 mm glass bottom dish using a 4-well insert with an adhesive biocompatible bottom surface (ibidi, 80466/80469), which was placed on top of the filter and filled with HyClone DMEM/High Modified media.

#### Transfection

Cells were transfected transiently with the plasmids pEGFP-C2_SEPT9_i1 (addgene plasmid #71609)^47^ or LAP2 Full I pAcGFP-N1 monomeric GFP (addgene plasmid #62044; gift from Dr. Eric Schirmer)^104^ using Lipofectamine 2000 transfection reagent (Invitrogen, 11668019) according to the manufacturer’s instructions with minor optimization. Transfections were performed on 35 mm dishes using Opti-MEM™ I Reduced Serum Medium (Gibco, 31985070). For plasmid DNA transfection, DNA-lipid complexes were prepared immediately before use. For each dish, 1.5 µg of plasmid DNA was diluted in 120 µl of Opti-MEM™ I, and 4 µl of Lipofectamine 2000 was diluted in 120 µl of Opti-MEM™ I in a separate tube. After a 5-minute incubation at room temperature, the diluted DNA and Lipofectamine 2000 were combined, gently mixed, and incubated for 20 minutes at room temperature. Immediately before adding the complexes, the culture medium was replaced with fresh DMEM High Glucose medium without antibiotics. The DNA–Lipofectamine 2000 complexes were then added dropwise to the cells and mixed by gently swirling the dishes. Cells were incubated with the complexes at 37 °C in a humidified atmosphere containing 5% CO₂ for 6 hours, after which the medium was replaced with complete DMEM.

### Generation of stable CRISPR/Cas9 genome-edited cells

To generate cell lines with stable depletion of SEPT9_i1, a guide RNA (gRNA) targeting the N-terminal segment of SEPT9 isoform 1 was designed within exon 2. The gRNA oligonucleotides with forward primary sequence 5’-CACCGCACGCGGACCTCCAGTGGC-3’ (Sept9V1-F2) and reverse primer sequence 5’-AAACGCCACTGGAGGTCCGCGTGC-3’ (Sept9V1-R2) were annealed and cloned into the plasmid backbone of the pSpCas9(BB)-2A-Puro (PX459) vector (Addgene, 48139), which expresses a Cas9/gRNA cassette, following standard cloning procedures. Correct insertion and sequence integrity were confirmed by sequencing.

U2OS and MDA-MB-231 cells were maintained under standard culture conditions (37 °C, 5% CO₂) in appropriate complete growth medium as described in the cell culture section. For transfection, cells were seeded one day prior to transfection to reach approximately 70–80% confluence and were then transfected with the pSpCas9(BB)-SEPT9_i1 gRNA plasmid using Lipofectamine 2000 according to the manufacturer’s instructions, with plasmid DNA and transfection reagent amounts optimized for each cell line. After 4–6 h, the transfection medium was replaced with fresh complete medium, and cells were allowed to recover overnight.

To enrich for transfected cells and facilitate stable line generation, transfected cultures were subjected to selection with puromycin (1 μg/ml) to eliminate non-transfected cells within 48 h. Following two days of puromycin selection, surviving cells were trypsinized and reseeded at very low density into 10 cm dishes to allow the formation of individual colonies derived from single cells. Colonies were manually isolated using a sterile filter paper with trypsin and transferred individually into the wells of a 48-well plate, where they were cultured for approximately two weeks with changes of media every 2–3 days. Clonal populations were expanded stepwise into larger dishes. Once sufficient cell numbers were obtained from each clone, whole-cell lysates were prepared and SEPT9_i1 protein levels were assessed by Western blot using an isoform-specific anti-SEPT9i1 antibody, with appropriate loading controls. SEPT9 exon 2 deletion was confirmed by genomic DNA sequencing using both the sequencing primers 5’-TAGTAGGGCTTGGGGGAGGATGTGTGTGAAGG-3’ (SeqCRISPseptV1F) and 5’-ACACACGGGCTTGCTCCCTGAGCCTGGGGGTG-3’ (SeqCRISPseptV1R).

### Confined cell migration and myosin hyperactivation assays

In the confined cell migration assays, we used the following Corning Costar Transwel plates: 24 mm Transwell with 3.0 µm pore polycarbonate membrane insert (Corning, 3452), 6.5 mm Transwell with 5.0 µm pore polycarbonate membrane insert (Corning, 3421), and 24 mm Transwell with 8.0 µm pore polycarbonate membrane insert (Corning, 3428). These were coated on both sides with Matrigel and laminin as described above. Two million cells (per 24 mm Transwell with 3.0 µm wide pores), one million cells (per 24 mm Transwell with 8.0 µm wide pores) and 0.5 million cells (per 6.5 mm Transwell with 5.0 µm pores) were resuspended in 1.5 ml serum-free medium and seeded directly onto the upper side of the Transwell membrane. The lower chamber received 2 ml of medium supplemented with 50 ng/ml EGF (Gibco, PHG0311) to establish a chemotactic gradient. Cells were allowed to migrate for 12 hours (8 µm wide pores), 24 hours (5 µm wide pores), or 36 hours (3 µm wide pores) at 37 °C and 5% CO₂.

After incubation, cells were washed twice with PBS to remove non-adherent cells and debris. Membranes were fixed in 4% paraformaldehyde (PFA) for 15 minutes at room temperature, quenched with 0.4% ammonium chloride for 10 minutes to reduce autofluorescence, and permeabilized as needed. Inserts were carefully removed from the Transwell plate, and membranes were mounted onto Parafilm for subsequent immunofluorescence staining.

For myosin II hyperactivation assays, cells were plated on collagen-coated glass coverslips or glass bottom dishes as described above, and cultured for 24 h prior to treatment with calyculin A (10 nM) or equal volume of DMSO (carrier control). Calyculin (Calyculin A from Discodermia calyx; In Solution, >98%) was purchased from Sigma-Aldrich (5.08226) and reconstituted with DMSO at stock concentration of 100 μM and kept at -20 °C after aliquoting. Cells were incubated with calyculin A for 15 or 30 minutes prior to fixation and permeabilization for imaging, or imaged live after 5 minutes of addition to glass bottom dishes.

### Cell viability assay

Cell viability following migration through the pores of the Transwell polycarbonate membranes was evaluated using a trypan blue exclusion assay (Fig. 5f). Briefly, two Transwell membranes (3 µm pore size) were used per sample replicate. A total of 2 × 10⁶ cells were seeded onto each Transwell membrane and stimulated with epidermal growth factor (EGF, 50 ng/mL) to promote directed migration. After 36 hours of incubation, cells that had traversed the membranes were recovered from the lower chamber by trypsinization, and collected in culture medium. Cells from the two membranes were pooled and centrifuged at 2,000 rpm. The resulting cell pellet was gently resuspended in 30 µL of PBS and subsequently diluted 1:1 (v/v) with 0.4% trypan blue solution (Invitrogen) for viability assessment. Stained cell suspensions were immediately loaded onto disposable Countess cell counting slides and analyzed using the Countess 3 Automated Cell Counter (Thermo Fisher Scientific).

The instrument showed reproducible image-based discrimination of viable (unstained) and non-viable (blue-stained) cells. For each condition, at least three independent biological replicates were quantified, and each replicate measurement represented the mean of three technical readings. The percentage of dead cells was calculated as Cell death (%) = (Number of trypan blue–positive cells/Total number of cells counted)×100.

### Immunofluorescence

#### Permeabilization and Fixation

Cells were cultured on the collagen-coated coverslips and subsequently fixed with 4% paraformaldehyde (Electron Microscopy Sciences, Cat # 15710) in phosphate-buffered saline (PBS) for 15 minutes at 37 °C following two PBS washes. To quench residual aldehyde groups, cells were incubated with 0.4% ammonium chloride (Sigma, catalog number A9434) for 5 minutes at room temperature. Permeabilization was performed using 0.2% Triton X-100 (Sigma Cat # X100) in PBS for 20 minutes at room temperature.

After permeabilization, cells were blocked in blocking buffer containing 2% bovine serum albumin (BSA; Sigma, Cat # A9647), and 0.1-0.2% Triton X-100 in PBS for 30 minutes. The samples were washed twice with PBS, blocked in 2% BSA in PBS for 30 minutes, and incubated with primary antibodies diluted in blocking buffer overnight at 4 °C. After primary antibody incubation, coverslips were washed and then incubated with appropriate fluorescently labeled secondary antibodies for 1 hour at room temperature in the dark.

Following immunostaining, coverslips were washed in PBS, stained with DAPI as necessary, and mounted onto frosted microscope slides (Globe Scientific, catalog number 1304W) using FluorSave™ reagent (Millipore, 345789). The slides were stored at 4 °C and protected from light until imaging.

#### Antibodies and fluorescent probes

The following antibodies were used for immunofluorescence microscopy and western blots: rabbit anti-Septin-9 (Proteintech, 10769), rabbit anti-SEPT7 (IBL America, 18991), rat anti-SEPT9_v1 4D2A5 (Cell Engineering Corporation, CEC-048), mouse anti-paxillin (BD Transduction Laboratories, 610051; clone 349), mouse monoclonal anti-phospho-Histone H2A.X (Ser139) clone JBW301 (MilliporeSigma, 05-636-I), mouse anti-Nuclear Pore Complex clone 414 (MilliporeSigma, N8786), mouse anti-paxillin (ND Transduction Laboratory, 61005), rabbit anti-zyxin (Proteintech, 10330), mouse anti-myosin IIA (MilliporeSigma, M8064), rabbit anti-phospho-Myosin Light Chain 2 (Thr18/Ser19) (Cell Signalling, 3674), and rabbit anti-vimentin (Proteintech, 10366), and mouse anti-GAPDH (Novus, Nb600-52SS).

The following fluorescent antibodies were used for immunodetection of primary antibodies in immunofluorescence experiments: Alexa Fluor 488 AffiniPure donkey anti-rat IgG (H+L) (Jackson ImmunoResearch, 712-545-153), Alexa Fluor 488 AffiniPure F(ab)₂ donkey anti-rabbit IgG (H+L) (Jackson ImmunoResearch, 712-545-153), AlexaFluor 488 AffiniPure F(ab)₂ donkey anti-mouse IgG (H+L) (Jackson ImmunoResearch, 715-546-150), Alexa Fluor 547 AffiniPure F(ab)₂ donkey anti-Rabbit IgG (H+L) (Jackson ImmunoResearch, 711-546-152).

Actin was labeled in fixed cells with AF488 Phalloidin (AAT Bioquest, 23153) and Phalloidin-iFluor 647 (Abcam, ab176759), and in live cells with SPY555-Actin (Cytoskeleton, CY-SC202), SPY555-FastAct (Cytoskeleton, CY-SC205) or SPY650-FastAct_X (Cytoskeleton, CY-SC502). Cell nuclei were stained with DAPI (1 µg/ml; ThermoFisher Scientific) in fixed cells and the Hoechst 3342 (Invitrogen) in live cells.

For western blots, the following dye–conjugated secondary antibodies were used: goat anti-rate IRDye680 RD (Li-Cor Biosciences, 925-68076), donkey anti-rabbit IRDye800 CW (Li-Cor Biosciences, 926-32213) and donkey anti-mouse IRDye680 (Li-Cor Biosciences, 926-32222).

### Microscopy and image processing

#### Super-resolution spinning disk confocal microscopy

Live cell imaging and imaging of fixed cells in confined migration assays was performed with a NIKON Ti2E motorized microscope with a super-resolution W1-SoRa Yokogawa spinning disk system equipped with a Hamamatsu ORCA-Fusion BT back-thinned camera, and an environmentally controlled enclosure (Okolab) for adjustment and maintenance of temperature, humidity, and CO2 during live-cell imaging. Images were acquired with a CFI60 PLAN APOCHROMAT LAMBDA D 60X Oil, N.A. 1.42, DIC and a SOLA Light Engine Gen III with 350-680nm unfiltered light output and full complement of excitation/emission filter sets, and a laser illumination LUN-F-XL output with 405 nm (50 mW), 488 nm (60 mW), 561 (50 mW) and 640 (40 mW) laser lines.

Following identification of cell regions of interest using the DAPI fluorescence, super-resolution imaging was conducted by setting Observation Mode to Confocal SoRa with a digital 2.5X magnification and an axial step size of 0.1–0.2 µm (1152 × 1152 pixels per slice, 12-bit depth, pinhole 1 AU) for acquisition of a z-series stack, which ensured high-resolution sampling along the Z-axis and accurate three-dimensional reconstruction of the cellular structures within the pores. Laser power was set to 10% to minimize photobleaching, with dwell time of 1.6 µs per pixel. Images were acquired using Nikon NIS-Elements C software (v. 6.4, Nikon Instruments).

Raw z-stacks were processed in Nikon NIS-Elements Advanced Research software (v. 6.4). A Gaussian noise reduction filter (σ = 1.0) was applied first to minimize background signal and enhance signal-to-noise ratio, followed by blind deconvolution (10 iterations, Richardson–Lucy algorithm) to enhance resolution and contrast. Processed stacks underwent 3D volume rendering (isosurface projection, threshold 20–50% max intensity) for pore-cell reconstruction, enabling detailed visualization and structural assessment of cells occupying or migrating through the membrane pores.

#### Wide-filed deconvolution microscopy

Imaging of fixed cells on 2D collagen-coated coverslips was performed with a NIKON Ti2E motorized widefield/deconvolution microscope system equipped with a Hamamatsu ORCA-Fusion BT back-thinned camera, Plan Apochromat Lambda D 60x oil immersion objective lens (N.A. 1.42) with phase contrast and DIC condensers and prisms, and a SPECTRA III illumination source for excitation at 390 nm, 440 nm, 475 nm, 510 nm, 555 nm, 575 nm, 637 nm and 748 nm wavelengths. band pass filters. Images were acquired with the Nikon Elements software with post-processing (denoising; LIM 2D/3D deconvolution) and analysis tool on a HP Z4 advanced imaging workstation. Samples were illuminated under conditions optimized to prevent photobleaching while ensuring adequate signal-to-noise ratio. Image stacks were acquired with 0.1–0.2 µm z-step intervals to capture the full three-dimensional structure of the specimen. Images were denoised and deconvolved, as indicated, using the Nikon Elements software.

### Western blots of cell extracts

MDA-MB-231, U2OS, and their respective SEPT9-deficient cell lines were cultured in 60 mm tissue culture dishes at a density of 1 × 10⁶ cells per plate for 24 hours under standard growth conditions. Following incubation, cells were washed once with ice-cold phosphate-buffered saline (PBS) and lysed on ice for 30 minutes in lysis buffer containing 150 mM NaCl, 50 mM Tris-HCl (pH 8.0), 20 mM NaF, and 1% NP-40, supplemented with Complete Mini EDTA-free protease inhibitor cocktail (Roche, Cat. No. 11836170001). Lysates were mixed gently by pipetting to ensure homogeneity and then subjected to two cycles of sonication (10 seconds on, followed by 5 minutes off) to disrupt cellular membranes and shear genomic DNA. After sonication, samples were centrifuged at 12,000 rpm for 15 minutes at 4 °C, and the clarified supernatants were collected as total protein extracts. Protein concentration was determined using a Pierce BCA protein assay kit (BioRad) and equal amounts of protein were mixed with 5X Laemmli sample buffer and boiled for 5 minutes at 95 °C to denature the proteins.

Samples were separated by SDS–polyacrylamide gel electrophoresis (SDS–PAGE) and subsequently transferred to nitrocellulose membranes using a wet transfer apparatus. Membranes were blocked for 1 hour at room temperature in blocking buffer (2% nonfat dry milk in TBS-T: 20 mM Tris-HCl, 150 mM NaCl, 0.1% Tween-20) to prevent nonspecific binding.

The membranes were incubated overnight at 4 °C with primary antibodies diluted in blocking buffer, followed by incubation with appropriate infrared dye–conjugated secondary antibodies (Li-Cor Biosciences-IRDye 800CW and IRDye 680RD) for 1 hour at room temperature. After washing, protein bands were visualized and quantified using the Odyssey CLx-Infrared Imaging System (Li-Cor Biosciences) according to the manufacturer’s instructions.

### Image Analysis and Quantifications

#### Quantification of actin cable decoration with zyxin, pMLC2 and SEPT9, and contact with paxillin

Images were first subjected to a noise reduction procedure and subsequent deconvolution using NIS-Elements software (Nikon) to improve signal-to-noise ratio and spatial resolution prior to quantitative analysis.

Quantification of zyxin, pMLC2 and SEPT9 association with actin filaments was conducted following three-dimensional (3D) image reconstruction of cells using NIS Elements software (Nikon Instruments). Three-dimensional image reconstructions were then generated from SoRa z-stacks comprising 20–30 optical sections using the “Show 3D View” function, thereby enabling accurate visualization and measurement of filament geometry in three dimensions. Zyxin fluorescence signals colocalizing with actin cables were identified, excluding signals located at filament termini, and individual actin cables with zyxin fluorescence were scored as zyxin-positive. Similarly, actin cables decorated with pMLC2 and SEPT9 independently of extent and region of the localization on filamentous actin were scored as positive. Actin cables with end-on contacts and overlaps with paxillin were also recorded in 3D volumes of cell regions confined in pores. The number of zyxin, pMLC2 or SEPT9 positive cables, and paxillin capped actin cables per cell was recorded, and calculated as percentage of total number of actin cables per cell after recording the absolute number of the latter. Percentiles for each cell were imported into GraphPad Prism (GraphPad Software) for statistical analysis and graphing.

#### Quantification of actin filament number, length, orientation, severing, buckling and retraction rates

Quantification of actin filament number and length was done using the Nikon Elements tool "3D Measurement" from the "Measure" software function, which enabled the marking of each actin cable, and the lengths were measured using the "3D Length" function. Results per each confined region were compiled in Microsoft Exel, and imported into GraphPad Prism 6 for data analysis, graph generation and statistical analysis. Actin cable angles of deviation from the vertical apicobasal axis of the pore were measured using the "3D Angle" tool of the Nikon Elements software, and recorded for each actin cable of the confined cell region.

In images of MEF cells cultured in 2D, actin stress fibers were segmented using the auto thresholding function in Fiji/Image J. Broken actin filaments were identified as individual phalloidin-labeled actin stress fibers with at least one discontinuity or gap along their lengths. Buckled actin filaments were identified as phalloidin-labeled actin stress fibers with at least one bent or wavy segment. The number of broken actin and buckles filaments per cell was scored after blinding for experimental conditions. The percentage of broken and buckles actin filaments was calculated as (number of broken or buckled filaments per cells/total filaments per cells) ×100%. Data were compared to control values and statistical analyses were performed using GraphPad Prism.

In time-lapse images of MEF cells, quantification of actin stress fiber severing sites as septin-coated or septin-free sites was done by applying a minimum of 10 μm length of actin coverage or non-coverage by GFP-SEPT9_i1 signal with respect to the site of breakage, which was visualized by the SPY555-Actin fluorescence signal. Sites of severing that localized within a ≥ 10 μm-long actin segment coated with GFP-SEPT9_i1 were scored as septin-positive, and sites of severing that localized within a ≥ 10 μm-long actin segment free of GFP-SEPT9_i1 were scored as negative. Following a severing event, retraction rates were measured by generating a thin sliced ROI along ther axis of breakage. The average velocity was calculated from time-distance montages (kymographs) from the initial breakage point to stoppage point or until the stress fiber retracted past the field of view or became ambiguous.

#### Quantification of nuclear areas and nuclear envelope rupture

Nuclear area was quantified from fluorescence microscopy images using Fiji. In images with DAPI fluorescence, the nuclear channel was isolated using “Image > Color > Split Channels,” and subsequent analysis was performed on the corresponding grayscale nuclear image. To segment nuclei, a global threshold was applied (“Image > Adjust > Threshold”) to distinguish nuclei from background, choosing an intensity range that rendered each nucleus as a single continuous region. Thresholds were applied using the “Apply” command to generate a binary image, and in cases of touching nuclei, the “Process > Binary > Watershed” function was used to separate adjacent objects. Before measurement, the set of parameters was defined under “Analyze > Set Measurements” enabling Area. Nuclear areas were then obtained in in µm² areas with “Analyze > Analyze Particles”. The resulting data containing individual nuclear areas was copied to excel file and subsequently imported into GraphPad Prism for statistical analysis and graph generation.

To quantify nuclear rupture events, fluorescence microscopy images of cells stained with DAPI and the mouse mAb 414 against nucleoporins, which outlined the nuclear envelope, were analyzed using Fiji (ImageJ). Images were first imported as multi-channel stacks, and each channel was separated to independently evaluate the nuclear membrane continuity that surrounds DNA. Background subtraction and intensity normalization were applied to improve signal-to-noise ratio. Using the nucleoporin channel, the nuclear boundary was visualized and segmented using manual thresholding. Rupture was defined as a discontinuity or loss of membrane signal continuity surrounding the nuclear DAPI mass. The DAPI channel was used to confirm the presence of DNA protrusions extending beyond these discontinuities. For each image, nuclei were counted and categorized as ruptured or intact. Quantification was expressed as the percentage of ruptured nuclei relative to the total number of nuclei analyzed per field of view. Multiple fields were evaluated for each condition, ensuring statistical robustness. Data were compiled and analyzed using Fiji’s measurement tools and exported for statistical processing in GraphPad Prism.

#### Quantification of DNA damage, micronuclei and nuclear blebs

DNA damage image analysis was performed using FIJI/ImageJ. Individual nuclei were segmented based on DAPI staining, and cells exhibiting at least one large γH2AX focus with surface area larger than 0.4 μm^2^ were classified as γH2AX positive. The fraction of γH2AX-positive cells was calculated as: percentage of γH2AX-positive cells= (Number of γH2AX-positive nuclei/Total number of nuclei analyzed) ×100 Micronuclei were identified and quantified as DAPI-positive structures separated from the main nuclear mass, with a diameter less than one-third of the primary nucleus, and surrounded by a continuous NUP414-positive envelope. The percentage of cells containing micronuclei was calculated as: Micronuclei (%) = (Number of cells with micronuclei/Total number of cells)×100.

Nuclear blebbing analysis was done in cells stained with mAb 414 against nucleoporins and counterstained with DAPI. A cell was scored as positive for nuclear blebbing if the DAPI signal displayed an irregular contour with at least one protruding bleb, which emerged from and/or outlined by a discontinuous or broken nuclear rim as evident by the nucleoporin fluorescence signal. The percentage of cells exhibiting nuclear blebs was calculated per each region imaged (field of view) as: Nuclear bleb (%) = (Number of cells with nuclear blebs/Total number of cells)×100. At least 50 cells from independent biological repicates were quantified per condition.

### Analysis of clinical genomic data

#### Data sources and acquisition

TCGA molecular and clinical data were retrieved from cBioPortal using the R package **cBioPortalData**. TCGA studies were screened for the availability of discrete GISTIC copy-number profiles, tumor mutation burden (TMB) annotations, and corresponding clinical and survival data. For each eligible study, study-level clinical data, *SEPT9* discrete copy-number calls, and TMB values were downloaded via the cBioPortal API and stored locally for downstream harmonization.

#### Data harmonization and quality control

Study-level tables were harmonized into a unified clinicogenomic analysis dataset using a study-aware merge strategy. Copy-number alterations and TMB values were required to be unique per study and sample, and clinical annotations were used to map samples to patients. Overall survival in months was used as the primary endpoint, with one survival record retained per patient.

To prevent duplication across TCGA study variants, sample identifiers appearing in multiple studies were resolved by retaining a single canonical record per sample. Samples with unresolved conflicts or missing key annotations were excluded. The final analysis dataset contained one row per unique tumor sample with consistent copy-number, mutational, clinical, and survival annotations.

#### Overall survival analysis

Overall survival (OS) analyses were performed using the harmonized TCGA clinicogenomic dataset. Samples were required to have non-missing OS time, event status, and a valid tumor-type annotation. *SEPT9* copy-number status was defined using GISTIC discrete calls (−2 to +2). Tumors with *SEPT9* gain or amplification (GISTIC ≥ +1) were compared to copy-neutral tumors (GISTIC = 0); deletion states were excluded.

Kaplan-Meier survival curves were estimated and compared using log-rank tests. Pan-cancer Cox proportional hazards models were additionally fit with tumor-type stratification to account for baseline hazard differences across cancer types. Proportional hazards assumptions were evaluated using Schoenfeld residuals.

#### Tumor-type enrichment of *SEPT9* copy-number alterations

Tumor-type enrichment of *SEPT9* copy-number alterations was evaluated using two binary exposure definitions: (i) gain or amplification (GISTIC ≥ +1) and (ii) amplification-only (GISTIC = +2). For each tumor type with sufficient sample size (N ≥ 20), enrichment was assessed using Fisher’s exact test by comparing the frequency of the CNA state within the tumor type to all other tumor types combined. P values were adjusted for multiple testing using the Benjamini–Hochberg procedure. Effect sizes were summarized as odds ratios and log2-transformed odds ratios for visualization.

#### Copy number-expression association

To assess whether *SEPT9* copy-number alterations are associated with increased transcript abundance, matched copy-number and mRNA expression data were analyzed from the TCGA breast cancer cohort accessed via cBioPortal (Firehose Legacy). Tumors were stratified by SEPT9 GISTIC copy-number state (−2, −1, 0, +1, +2), and *SEPT9* mRNA expression values were extracted as provided by the portal. Differences in expression across copy-number strata were evaluated using a Kruskal-Wallis ANOVA test.

Where indicated, monotonic association between copy-number state and mRNA expression was additionally assessed using Spearman rank correlation, treating copy-number state as an ordinal variable.

#### Association between *SEPT9* copy number and chromosomal instability

Chromosomal instability was quantified using fractional genome altered (FGA), defined as the proportion of the genome affected by copy-number gains or losses. Analyses were restricted to tumors with non-missing FGA values and available *SEPT9* GISTIC copy-number calls.

Tumors were stratified by *SEPT9* copy-number status as gain or amplification (GISTIC ≥ +1) versus copy-neutral tumors. Differences in the distribution of FGA between groups were evaluated using non-parametric density comparisons.

#### Association of *SEPT9* copy number with tumor stage and grade

Tumor stage and histologic grade were obtained from harmonized TCGA clinical annotations and reflects American Joint Committee on Cancer (AJCC) staging criteria. AJCC stage integrates the extent of the primary tumor, regional lymph node involvement, and distant metastasis to classify disease severity into stages I–IV. For pan-cancer analyses, detailed AJCC sub-stages were collapsed into a simplified categorical variable to enable consistent comparisons across tumor types. Tumor grade information was derived from pathology-reported histologic grade annotations. Where available, grades were classified using conventional four-tier systems (G1-G4), reflecting increasing degrees of tumor dedifferentiation and proliferative aggressiveness, with G1 representing well-differentiated tumors and G4 representing poorly differentiated or anaplastic tumors. To ensure interpretability and consistency across cancer types, analyses of tumor grade were restricted to samples annotated as G1–G4, excluding ambiguous or non-standard categories (e.g., GX or unknown). *SEPT9* copy-number status was defined from discrete copy-number calls and categorized as copy-neutral, gain, or amplification. Associations between *SEPT9* copy-number state and clinical variables were evaluated using contingency table analyses. Differences in the distribution of tumor stage and tumor grade across *SEPT9* copy-number states were assessed using Pearson’s chi-square tests of independence. All analyses were performed in a pan-cancer cohort, and reported p-values are two-sided.

#### Tumor mutation burden and hypermutation analyses

Tumor mutation burden (TMB) was analyzed in the harmonized TCGA clinicogenomic dataset (see Source Data TumorMutationBurdenAnalysis). To avoid mixing incompatible TMB annotations, analyses were restricted to samples with non-missing numeric TMB values. *SEPT9* copy-number status was defined from GISTIC discrete calls, comparing tumors with gain or amplification (cna_discrete ≥ +1) to copy-neutral tumors (cna_discrete = 0); deletion states were excluded from these comparisons. To define hypermutation in a tumor-type-adjusted manner, high TMB was defined within each cancer type as tumors in the top 20% of the TMB distribution (80^th^ percentile cutoff), computed for tumor types with ≥20 evaluable samples. Enrichment of high-TMB tumors among *SEPT9*-altered versus copy-neutral tumors was evaluated using Fisher’s exact tests in pan-cancer analyses and within the subset of tumor types previously identified as enriched for *SEPT9* copy-number alteration. To account for tumor-type stratification, Cochran-Mantel-Haenszel tests were additionally performed with cancer type as the stratification variable.

For continuous TMB analyses, TMB was log-transformed as log10(TMB + 0.1). Associations between *SEPT9* status and continuous TMB were tested using linear models including tumor type as a fixed effect (log10(TMB + 0.1) ∼ *SEPT9* status + cancer type). As a tumor-normalized sensitivity analysis, within-tumor type z-scores of log10(TMB+0.1) were computed and modeled as a function of *SEPT9* status.

#### Statistical Analyses

Statistical analysis of data and graph generation was performed with GraphPad Prism statistical analysis software. Violin plots display the individual values of all data sets, and indicate the median (black line), first and third quartiles (colored lines). Data set groups were first checked for standard deviation (SD) variance using the Prism’s Descriptive Statistics tool and then, assessed for normal distribution of variance using the D’Agostino and Pearson and Shapiro-Wilk normality tests. For pair-wise comparisons of data with normal distributions, a student’s t test was used if SDs were equal, or a Welch’s t-test if SDs were unequal. The Mann-Whitney test was used for non-normally distributed data. For statistical analysis of multiple groups with normally distributed data, a Brown-Forsyth and Welch ANOVA test was performed with a post-hoc Dunnett T3 test for multiple pair-wise comparisons. For data that were not normally distributed, a non-parametric Kruskal-Wallis ANOVA test was performed with a post-hoc Dunn’s test for multiple comparisons. Data comparison in survival Kaplan-Meier graphs were done with logP or hazard ratio, and chi square analysis was used to assess comparisons between groups with categorical data. Additional statistical tests in the analysis of clinical genomic data were as indicated above.

## Supporting information

Supplementary Video 1: Actin cables surround the pore-confined nuclear mass and contact paxillin during 3D confined migration

Supplementary Video 2: Septins align vertically with the actin cables that surround the pore-confined nuclear mass

Supplementary Video 3: SEPT9 colocalizes with pMLC2 and vertical actin cables within the pore-confined cytoplasm.

Supplementary Video 4: Nuclear membrane and actin dynamics in a pore-confined cell region.

Supplementary Video 5: Perinuclear SEPT9 dynamics in a pore-confined cell region.

Supplementary Video 6: SEPT9-coated actin filament segments retract more slowly

Supplementary Video 7: Supplementary Video 7 Septin-associated actomyosin cables surround the nucleus in a pore-confined region of an MDA-MB-231 cell

## Acknowledgments

We are grateful to Drs. Ernst-Martin Füchtbauer (University of Aarhus, Denmark), Eric Schirmer (University of Edinburgh) and Greg Alushin (Rockefeller University) for cell and reagents. We thank the neighboring labs of our UVA colleagues Drs. Seham Ebrahim and Doug DeSimone for help with reagents, and the UVA Summer Research Internship Program (SRIP) and SRIP interns Aidee Hernandez (Amherst College) and Elizabeth Montgomery (Liberty University) for technical help. This work was supported by funding from the National Institute on Aging of the National Institutes of Health (NIH) grant R01 AG068908 to C.M.; Rutgers Cancer Institute funding to C.M. from the National Cancer Institute (NIH) grant P30CA072720-25; Breast Cancer Research funding within the University of Virginia Comprehensive Cancer Center (UVACCC) to E.T.S.; National Institute of General Medical Sciences (NIH) funding under the R35 award GM136337 to E.T.S.

## Supplementary Videos

**Supplementary Video 1**

**Actin cables surround the pore-confined nuclear mass and contact paxillin during 3D confined migration.** Super-resolution SoRa spinning disk confocal microscopy of a U2OS cell region confined within an 8 μm wide pore, showing F-actin (phalloidin, magenta), paxillin (green), and nucleus (DAPI, blue) after 3D volume rendering. The image rotates 180° degrees around the vertical axis of the pore. At the top of the rotating view, background fluorescence from actin filaments and the nucleus is visible from the cell region attached to the apical side of the porous Transwell membrane. The beaded appearance of some actin cables is due to sub-Nyquist sampling, which was necessary to offset photobleaching during acquisition of a large image z-stack. Image width, ∼10 μm.

**Supplementary Video 2**

**Septins align vertically with the actin cables that surround the pore-confined nuclear mass.**

Super-resolution SoRa spinning disk confocal microscopy of a U2OS cell region confined within an 8 μm wide pore, showing F-actin (phalloidin, magenta), SEPT9 (green) and nucleus (DAPI, blue) after 3D volume rendering. Image rotates 180° degrees around the vertical axis of the pore. The banded appearance of some actin cables is due to the sub-Nyquist sampling during the acquisition of a large image z-stack, which was applied to minimize photobleaching. Image width, ∼8 μm.

**Supplementary Video 3**

**SEPT9 colocalizes with pMLC2 and vertical actin cables within the pore-confined cytoplasm.**

Super-resolution SoRa spinning disk confocal microscopy of a U2OS cell region confined within an 8 μm wide pore, showing F-actin (phalloidin, magenta), SEPT9 (grayscale) and pMLC2 (green) after 3D volume rendering. The fluorescence signals are sequentially displayed to demonstrate colocalization: pMLC signal fades first, followed by SEPT9, revealing actin cables alone (magenta). Subsequently, pMLC2 and SEPT9 signals are sequentially restored, after which the pMLC2 and actin signals are removed to show SEPT9 alone. Gradual merging of the pMLC2 and F-actin signals with SEPT9 demonstrates colocalization of these proteins. Scale bar, ∼2 μm.

**Supplementary Video 4**

**Nuclear membrane and actin dynamics in a pore-confined cell region.** Time-lapse 3D super-resolution SoRa spinning disk confocal microscopy of a U2OS cell region confined within an 8 μm wide pore, epxressing LAP2β-AcGFP-1 (green, nuclear membrane) and labeled with SPY555-FastAct (magenta, F-actin). Three-dimensional image stacks were acquired every 10 minutes for 6 h, and rendered in 3D. Left panel shows LAP2β-AcGFP-1 alone; right panel shows merged channels. Actin-labeled structures emerge after 2 h. Image width, 8 μm.

**Supplementary Video 5**

**Perinuclear SEPT9 dynamics in a pore-confined cell region.** Time-lapse 4D super-resolution SoRa spinning disk confocal microscopy of a U2OS cell region confined within an 8 μm wide pore, expressing GFP-SEPT9_i1 (green) and labeled with Hoechst (blue; nucleus). Three-dimensional image stacks were acquired at 10-minute intervals for 190 minutes and rendered in 3D. Septin cables flank the nucleus vertically and fluctuate dynamically in position and length. Scale bar, 2 μm.

**Supplementary Video 6**

**SEPT9-coated actin filament segments retract more slowly than the septin-free segments after severing induced by myosin II hyper-contractility.** Time-lapse spinning disk confocal microscopy of *SEPT9*^-/-^ MEFs expressing GFP-SEPT9_i1 (green) and labeled with SPY555-FastAct (magenta; F-actin). Cells were treated with calyculin A (10 nM) for 15 minutes prior to live imaging to induce myosin hyper-contractility and actin severing events. Arrows indicate the site of actin severing and track the retracting end of the septin-free segment, which retracts faster than its septin-coated counterpart. Images were captured at 15 second intervals. Scale bar, 4 μm.

**Supplementary Video 7**

**Septin-associated actomyosin cables surround the nucleus in a pore-confined region of an MDA-MB-231 cell.** Super-resolution SoRa spinning disk confocal microscopy of an MDA-MB-231 cell region confined within a 3 μm wide pore, showing F-actin (phalloidin, magenta), SEPT9 (grayscale) and pMLC2 (green) after 3D volume rendering. The image initially rotates 90° degrees around the horizontal xy axis for viewing in 3D sideways from an oblique angle. The fluorescence signals are sequentially displayed: pMLC2 and SEPT9 signals fade first simultaneously, revealing actin cables alone (magenta). Subsequently, SEPT9 signal is restored prior to fading, and overlaying of pMLC2 onto F-actin with a 180°C rotation around the vertical axis of the pore. The beaded appearance of some actin cables is due to sub-Nyquist sampling, which was necessary to offset photobleaching during acquisition of a large image z-stack. Scale bar, ∼2 μm.

---

**Extended Data, Fig. 1:**
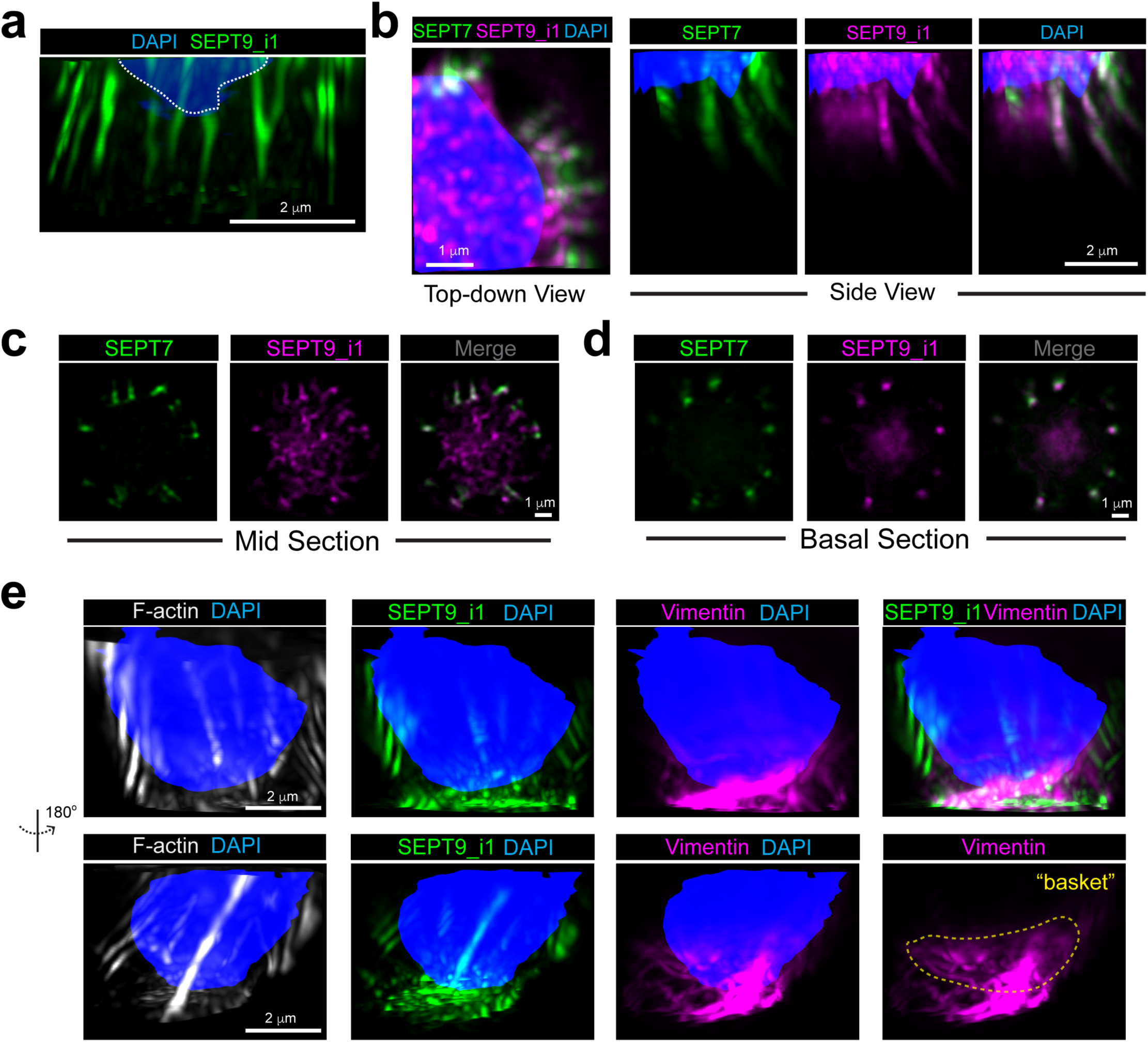
Perinuclear septin cables consist of SEPT9_i1 and SEPT7, and are spatially distinct from the nuclear vimentin basket. **a**, Sagittal view of SEPT9_i1 (green) and the nucleus (DAPI, blue) in a 3D volume region of a U2OS cell confined in an 8 mm-wide pore. Scale bar, 2 μm. **b,** Top-down and sagittal side views of a confined U2OS subcellular region after staining for SEPT7 (green), SEPT9_i1 (magenta) and the nucleus (DAPI, blue). Images were 3D reconstructed, and volume rendered. Scale bars, 1 μm (top-down view) and 2 μm (sideview). **c-d,** Single optical sections of a confined U2OS cell region taken from medial (c) and basal (d) depths after staining for SEPT7 (green) and SEPT9_i1 (magenta). Scale bar, 1 μm. **e,** Sagittal sideviews of the nucleus (DAPI, blue), F-actin (phalloidin, grayscale), SEPT9_i1 (green) and vimentin (magenta) in a 3D volume region of a U2OS cell confined in an 8 mm-wide pore. The view shown in the top row of images is rotated 180 degrees in the bottom row. The perinuclear vimentin basket is outlined with a yellow dashed line. Scale bars, 2 μm. All U2OS cell images were acquired with super-resolution SoRa SDCM 12 h after plating on 8 μm-wide pores.

**Extended Data, Fig. 2:**
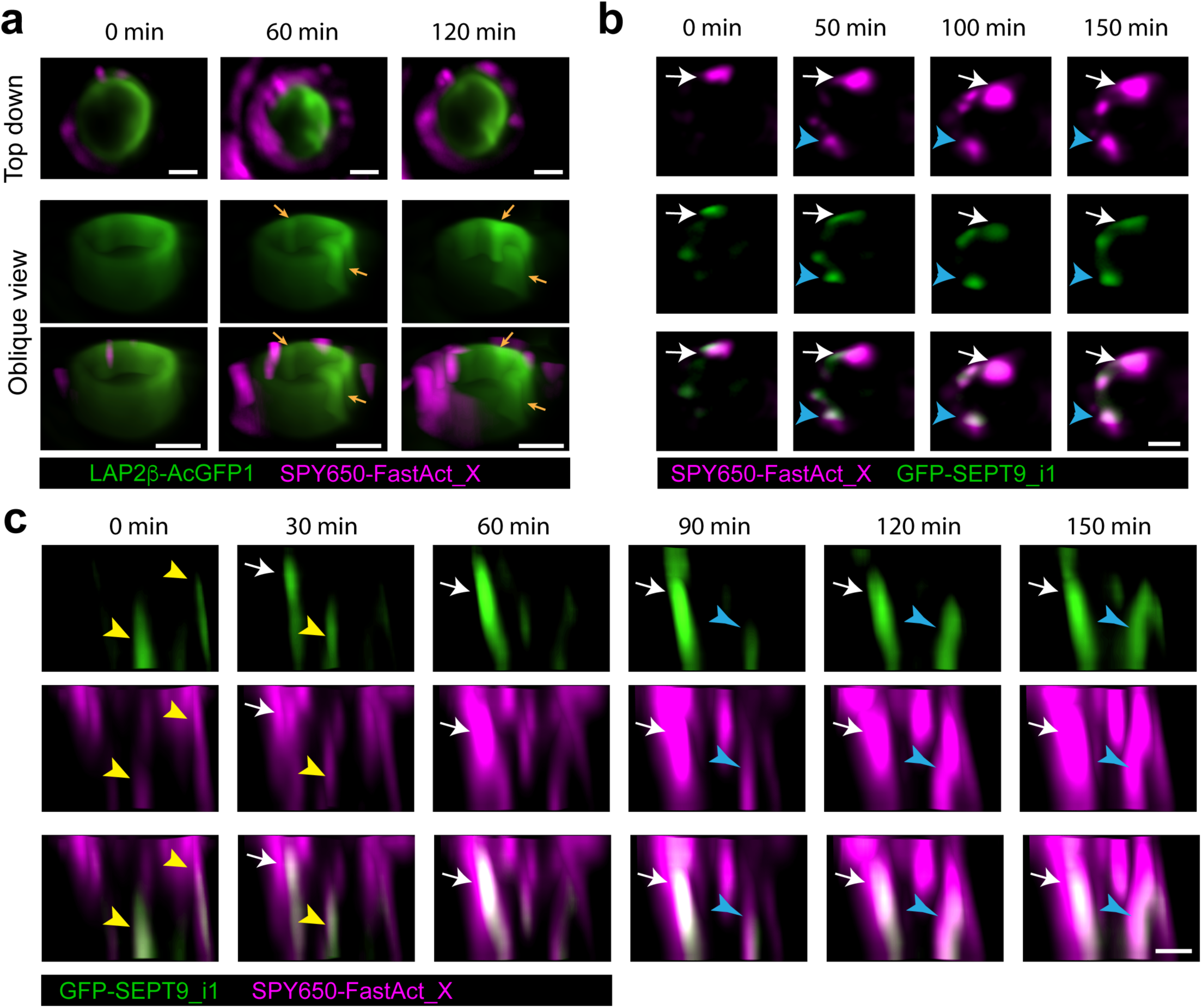
Spatiotemporal dynamics of perinuclear actin and septin cables in confined regions of cell migration. **a**, Top-down (top) and oblique angle (middle, bottom) views of still frames from time-lapse 3D SoRa SDCM imaging of LAP2β-AcGFP1 (green) and SPY555_FastAct_X (magenta) in the U2OS cell region confined in a Matrigel/laminin-filled 8 μm-wide pore. Images were rendered and reconstructed in 3D for each time point. Angled side views are more zoomed in on the nuclear envelope, which appears to surround the pore. Nuclear envelope creases and folds (arrows) are seen in the 60 and 120 minute time frames. Note that the nuclear envelope preserves its continuity in the 120 minute timeframe, but the front side becomes dim due to higher signal intensity in the creased regions of the 3D-rendered image. Scale bars, 2 μm. **b,** Time-lapse frames of a single optical slice from SoRa SDCM imaging of a U2OS cell region confined in a Matrigel/laminin-filled 8 μm-wide pore. U2OS cells were transfected with GFP-SEPT9_i1 (green) and labeled live with SPY650_FastAct_X (magents). White arrows point to actin that colocalizes with GFP-SEPT9_i1, and enlarges overtime. Blue arrows point to actoseptin that appears in frame de novo and remains spatially stable overtime. Scale bars, 2 μm. **c,** Still frames show side views of 3D volume images from time-lapse 3D SoRa SDCM imaging of GFP-SEPT9_i1 (green) and SPY650_FastAct_X (magenta) in a U2OS cell region confined in a Matrigel/laminin-filled 8 μm-wide pore. Yellow arrowheads point to short-lived actoseptin cables. White arrows point to longer lived actoseptin cables which become brighter and thicker. Blue arrows point to de novo generation of actoseptin cables that grow longitudinally and laterally. Scale bar, 1 μm.

**Extended Data, Fig. 3:**
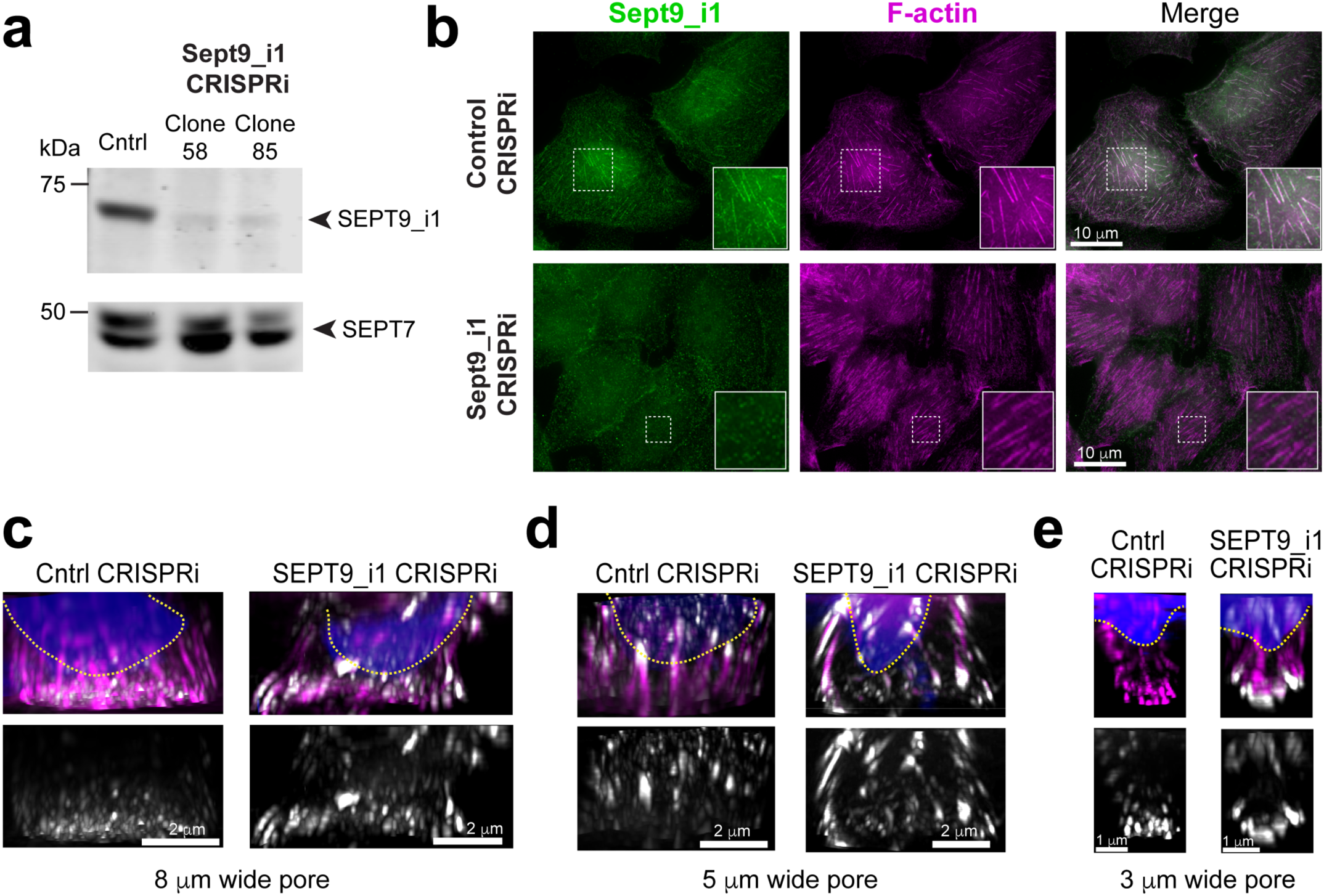
Septin expression in genome edited U2OS cells, and zyxin increase in confined SEPT9_i1-depleted U2OS cells. **a**, Western blots of lysates of parental U2OS and cell clones after selection for stable expression of SEPT9_i1 targeting sgRNA relative. Cell lysates were blotted with antibodies against the SEPT9_i1 isoform and SEPT7. **b,** Wide-field microscopy images of U2OS cells that stably express control and SEPT9_i1 sgRNAs after staining for F-actin (phalloidin, magenta) and the SEPT9_i1 isoform (green). Insets show outlined regions in higher magnification. Scale bars, 20 μm. **c-e,** Sagittal side views of 3D rendered images of the confined regions of control and SEPT9_i1-depleted U2OS cells in 8 μm (c), 5 μm (d) and 3 μm (e) wide pores. Cells were stained with phalloidin (F-actin, magenta) and antibody against zyxin (grayscale). Nuclear areas are outlined with dashed yellow line. Outlined regions with zyxin-decorated actin cables are shown in higher magnification. Scale bars, 2 μm (c, d) and 1 μm (e).

**Extended Data, Fig. 4:**
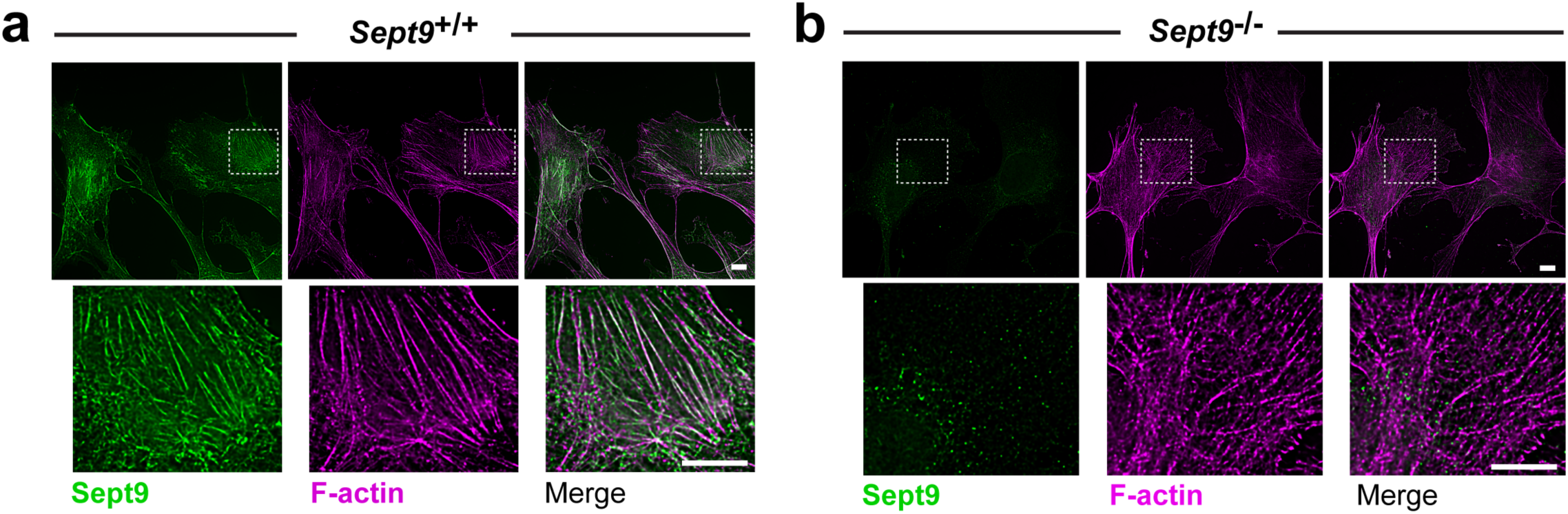
SEPT9 expression and localization in *Sept9*^+/+^ and *Sept9*^-/-^ MEFs. **a**, Wide-field microscopy images of *Sept9*^+/+^ MEFs on collagen type I ECM after staining for F-actin (phalloidin, magenta) and endogenous Sept9 with an antibody against all isoforms. Bottom panels show in higher magnification the outlined region with Sept9 filaments colocalizing with actin stress fibers. Scale bars, 10 μm. **b,** Wide-field microscopy images of *Sept9*^-/-^ MEFs on collagen type I ECM after staining for F-actin (phalloidin, magenta) and endogenous Sept9 (green) with an antibody against all isoforms. Bottom panels show in higher magnification an outlined region with actin stress fibers which lack Sept9 filaments. Absence of Sept9 expression is evident by the faint spotty background-level fluorescence signal. Scale bars, 10 μm.

**Extended Data, Fig. 5:**
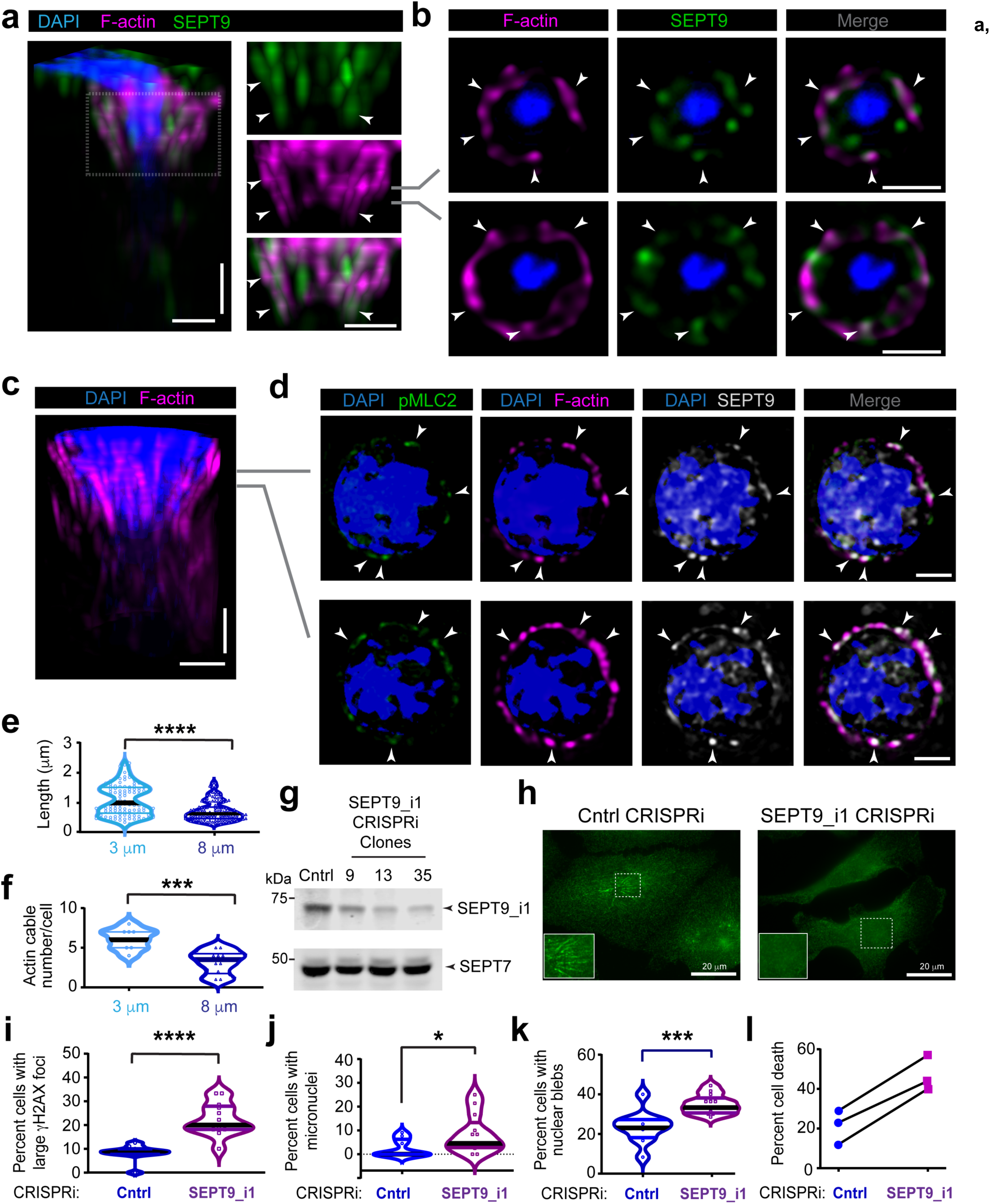
SEPT9 expression and localization in MDA-MB-231 cells during confined migration, and effects of SEPT9_i1 depletion on genomic integrity and cell viability. **a**, Sagittal 3D volume view of an MDA-MB-231 cell region confined in a Matrigel-filled 3 μm wide pore after staining for nucleus (DAPI, blue), F-actin (phalloidin, magenta) and SEPT9 (green). Images were acquired with 3D SoRa SDCM and rendered in 3D. Outlined regions is shown in higher magnification. Scale bars, 1 μm. **b,** Images show single SoRa SDCM optical slices taken from the indicated depths of the 3 μm wide pore (see lines in **a**). Arrowheads point to F-actin and SEPT9 colocalization. Scale bars, 1 μm. **c,** Sagittal 3D volume view of an MDA-MB-231 cell region confined in a Matrigel-filled 3 μm wide pore after staining for DAPI (blue), F-actin (phalloidin, magenta). Images were acquired with 3D SoRa SDCM and rendered in 3D. Scale bars, 1 μm. **d,** Images show single SoRa SDCM optical slices taken from the indicated depths of the 3 μm wide pore (see lines in **c**), and stained for nucleus (DAPI, blue), F-actin (phalloidin, magenta), pMLC2 (green) and SEPT9 (grayscale). Arrowheads point to F-actin, SEPT9 and pMLC2 colocalization. Scale bars, 1 μm. **e,** Violin plots of actin cable lengths in MDA-MB-231 cell region confined in Matrigel-filled 3 μm (*n =* 106 filaments, 9 cells) and 8 μm (*n =* 131, 10 cells) pores. Statistical comparison was done with a Mann Whitney U-test. ****, p<0.0001. **f,** Violin plots of the number of actin cables with length >0.5 μm per MDA-MB-231 cell region confined in Matrigel/laminin-filled 3 μm (*n* = 9 cells) and 8 μm (*n =* 10 cells) pores. Statistical comparison was done with an unpaired student t-test. ***, p<0.001. **g,** Western blots of lysates of MDA-MB-231 cell clones 9, 13 (used below in panels **i-l),** 58 (used in Fig. 5a**-f**) after selection for stable expression of parental control and SEPT9_i1 targeting sgRNAs. Cell lysates were blotted with antibodies against the SEPT9_i1 isoform and SEPT7. **h,** Wide-field microscopy images of MDA-MB-231 cells (clone 58) that stably express control and SEPT9_i1 sgRNAs after immunostaining with an isoform-specific antibody against endogenous SEPT9_i1 green). Insets show outlined regions in higher magnification. **i,** Violin plots show percentage of cells with large γH2AX foci after imaging basal regions of membranes with 3 μm wide pores containing parental control (*n* = 11 regions; 121 cells) and SEPT9_i1-depleted MDA-MB-231 cells (clone 13; *n* = 13; 147 cells) with large γH2AX foci. Statistical comparison was done with the Mann-Whitney U-test. ****, p<0.0001. **j,** Violin plots show percentage of cells with micronuclei after imaging basal regions of polycarbonate membranes with 3 μm wide pores containing parental control (*n* = 11 regions; 121 cells) and SEPT9_i1-depleted MDA-MB-231 cells (clone 13; *n* = 13 regions; 147 cells) with micronuclei. Statistical comparison was done with the Mann-Whitney U-test. *, p<0.05. **k,** Violin plots show percentage of cells with nuclear blebs after imaging basal regions of polycarbonate membranes with 3 μm wide pores containing parental control (*n* = 11 regions; 121 cells) and SEPT9_i1-depleted MDA-MB-231 cells (clone 13; *n* = 13 regions; 147 cells) with nuclear blebs. Statistical comparison was done with an unpaired t-test with Welch correction. ***, p<0.001. **l,** Percentage of Trypan blue-positive cells after trypsinization of parental control (*n* = 4185, 3400, 3660) and SEPT9_i1-depleted MDA-MB-231 cells (*n* = 5500, 2620, 4710) from the basal surfaces of Transwell polycarbonate membranes with 3 μm wide pores, following 36 h of apicobasal transmigration. Results from three independent experiments are shown. All violin plots show median (black line), first and third quartiles (colored lines).

**Extended Data, Fig. 6.**
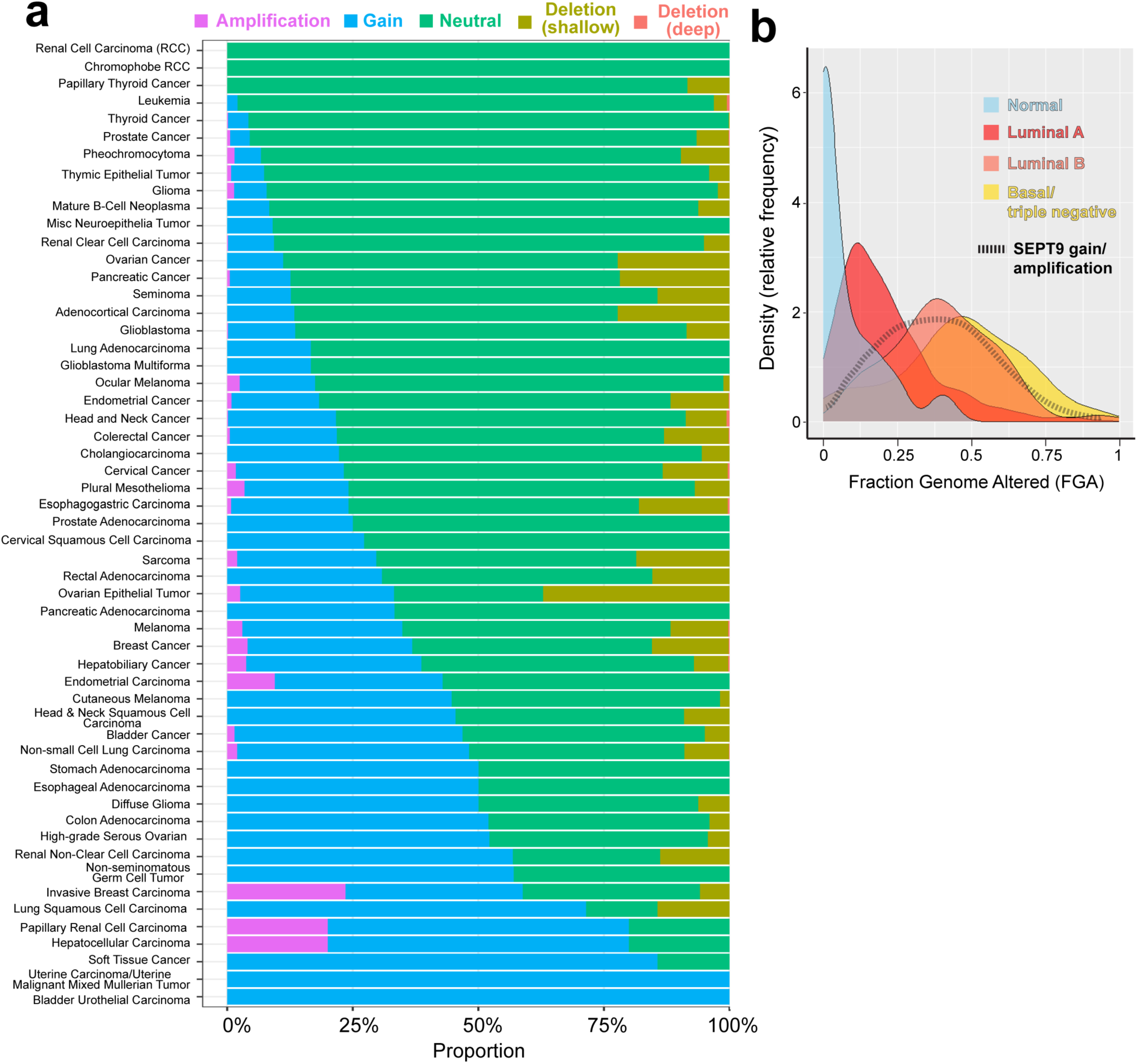
*SEPT9* gene copy amplification associates with thresholds of lower genomic alteration in aggressive tumors. **a**, Stacked bar plots showing the proportions of each indicated tumor type harboring *SEPT9* copy-number amplification (magenta), gain (cyan), shallow (olive) and deep depletion (pink), or copy-neutral status (green). Data are shown for pan-cancer primary tumors from TCGA (*n* = 11,245). **b,** Kernel density plots showing the distribution of FGA values in normal breast tissue (*n* = 36), luminal A breast cancer (*n* = 496), luminal B breast cancer (*n* = 197), basal/triple negative breast cancer (*n* = 171) and all breast tumors harboring *SEPT9* copy-number gain or amplification (*n* = 403).

